# Low nitrogen availability inhibits the phosphorus starvation response in maize (*Zea mays* ssp. *mays* L.)

**DOI:** 10.1101/2020.10.29.360487

**Authors:** J. Vladimir Torres-Rodríguez, M. Nancy Salazar-Vidal, Ricardo A. Chávez Montes, Julio A. Massange-Sánchez, C. Stewart Gillmor, Ruairidh J. H. Sawers

## Abstract

**Background:** Nitrogen (N) and phosphorus (P) are macronutrients essential for crop growth and productivity. In cultivated fields, N and P levels are rarely sufficient, contributing to the yield gap between realized and potential production. Fertilizer application increases nutrient availability, but not all farmers have access to fertilizers, nor are current rates of application sustainable or environmentally desirable. Transcriptomic studies of cereal crops have revealed dramatic responses to either low N or low P single stress treatments. In the field, however, levels of both N and P may be suboptimal. The interaction between N and P starvation responses remains to be fully characterized.

**Results:** We characterized growth and root and leaf transcriptomes of young maize plants under nutrient replete, low N, low P or combined low NP conditions. We identified 1,555 genes to respond to our nutrient treatments, in one or both tissues. A large group of genes, including many classical P starvation response genes, were regulated antagonistically between low N and P conditions. An additional experiment over a range of N availability indicated that a mild reduction in N levels was sufficient to repress the low P induction of P starvation genes. Although expression of P transporter genes was repressed under low N or low NP, we confirmed earlier reports of P hyper accumulation under N limitation.

**Conclusions:** Transcriptional responses to low N or P were distinct, with few genes responding in a similar way to the two single stress treatments. In combined NP stress, the low N response dominated, and the P starvation response was largely suppressed. A reduction in N availability was sufficient to repress the induction of P starvation associated genes. We conclude that activation of the transcriptional response to P starvation in maize is contingent on sufficient N availability.

## BACKGROUND

Nitrogen (N) and Phosphorus (P) are essential macronutrients required for multiple biological processes [1–5]. N is a component of all proteins and of the chlorophyll required for photosynthetic carbon fixation. P is required to produce the phospholipids that constitute the membranes that surround cells and intracellular organelles. Furthermore, N and P are both structural components of nucleic acids, including the abundant RNA molecules that play a key role in protein synthesis. The demand for these macronutrients is such that N and P availability in agricultural soils is rarely sufficient to realize the full yield potential of crops [6, 7]. P reacts readily with other elements, such as aluminum in acid soils or calcium in alkaline soils, leaving it immobile in the upper layers of the soil, reducing its availability to plants [8, 9]. By contrast, N, largely present in the form of nitrate, is mobile and tends to move to deeper soil layers where it may be beyond the reach of plant root systems [10]. In low-input systems, N and P deficiencies remain a major constraint on productivity. In high-input systems, the problem of N and P limitation is mitigated by chemical fertilizer addition, although current levels of application are neither sustainable nor desirable given associated environmental impacts [11]. Industrial N fixation is energetically costly and contributes to greenhouse gas production [12]. High grade phosphate rock is a non-renewable resource, predicted to be exhausted before the end of this century [13]. For these reasons, increasing N and P efficiency has been identified as a major target for plant breeding and agricultural management [11, 14].

Studies under controlled conditions, largely in the model plants *Arabidopsis thaliana* and rice (*Oryza sativa*), have identified physiological and developmental responses to low N or P stress, coupled with underlying large-scale changes in gene expression (the N starvation response or NSR, and P starvation response or PSR, respectively; [15–20]). A common response to nutrient deficiency is to promote uptake by increasing the abundance of high-affinity transporter proteins in the roots. Under N limitation, genes encoding the NTR2 nitrate transporters (working at N concentrations < 250 μM) are induced [21–23]. Similarly, genes encoding PHT1 P transporters (working at P concentration < 10 μM) are induced by P limitation [24–27]. Further aspects of the NSR include the downregulation of genes associated with nitrate assimilation, amino acid, oligosaccharide and nucleic acid biosynthesis [15, 28]. The PSR includes the induction of purple acid phosphatases (PAPs) involved in recycling internal and external P from organic pools, altered polysaccharide metabolism, and remodeling of lipid membranes to reduce the requirement for phospholipids [29–31]. Interestingly, aspects of the N and P responses appear to be antagonistic [15, 30, 32, 33]. Under single N limitation, many genes induced as part of the PSR are repressed. While molecular responses may be defined with respect to one nutrient at a time, it has long been appreciated that a deficiency in one element can impact the response to a second element, and that the effects of different nutrient deficiencies are not necessarily additive [34–40]. Thus, it is difficult to predict the transcriptomic response to a combination of N and P deficiency from the single stress data – especially in the context of antagonistically regulated genes. Several studies, however, have now demonstrated clear points of molecular interaction between N and P signaling pathways.

One of the first molecular links between N and P signaling was the identification of the SPX-RING (SPX domain: named after the Suppressor of Yeast gpa1, the yeast Phosphatase 81 and the human Xenotropic and Polytropic Retrovirus receptor 1; RING domain: Really Interesting New Gene) protein NITROGEN LIMITATION ADAPTATION (NLA1) in *Arabidopsis*. *Atnla1* mutants fail to adapt to low N conditions and exhibit early senescence [41] associated with P toxicity [42]. Further studies have shown that AtNLA directly targets PHT1 transporters for degradation in a N-dependent manner [43] as well as targeting the NRT1.7 nitrate transporter [44]. Under P starvation, downregulation of *AtNLA* by the P starvation inducible microRNA miR827 [42] promotes accumulation of PHT1. Rice OsNLA also regulates PHT1 abundance [45] and modulates P accumulation in an N-dependent manner [46]. However, in rice, miR827 does not target *OsNLA*, nor do N and P levels regulate *OsNLA* transcript accumulation, revealing regulatory differences compared to *Arabidopsis* [45, 47].

The MYB-CC transcription factor AtPHR1 plays a central role in activating the PSR [48]. Under high P, OsPHR2, the rice ortholog of AtPHR1, is sequestered by the SPX protein OsSPX4 preventing its translocation into the nucleus and activation of PSR genes [49]. Under P starvation, the 26S proteasome degrades OsSPX4, allowing OsPHR2 to activate its targets. Recently, the N-regulated OsNRT1.1b nitrate transporter has been shown to be required for OsSPX4 degradation. Under N starvation, levels of OsNRT1.1b are reduced, freeing OsSPX4 from turnover and leading to inhibition of the PSR [50]. Interestingly, OsSPX4 not only sequesters OsPHR2, but also the NIN-like protein OsNLP3, a central regulator of nitrogen response in rice [50]. These studies, and others, have demonstrated the interaction of N and P responses, and identified the SPX domain containing proteins as playing an important role in their coordination.

Maize is one of the world’s most economically important crops. Limitation of N or P represents a significant constraint on maize productivity worldwide [51–54]. Work in *Arabidopsis* and rice has begun to define the interactions between N and P signaling networks. Nevertheless, much remains to be discovered before we can apply this knowledge to the design of more efficient management practices or the development of more nutrient efficient crop varieties. Here, we report whole transcriptome data for the leaves and roots of maize seedlings grown in nutrient replete, low N, low P and a combined low NP stress. We observed antagonism between responses to single low N and low P treatments, with the low N response dominating in the combined low NP treatment. We further show that even a mild reduction in N availability is sufficient to suppress components of the maize PSR.

## RESULTS

### Growth of maize seedlings was reduced under low N and P treatments

To select material in which to characterize transcriptional responses to combined N and P limitation, we first characterized the growth of maize plants grown for 40 days after emergence under complete nutrient conditions (Full), reduced N (LowN: 9% of complete concentration), reduced P (LowP: 3% of complete concentration), and under combined reduced N and P (LowNP). Plants were grown in 1m tall, 15cm diameter (∼17L volume) PVC tubes, providing sufficient depth for root development (Fig. 1A). We followed plant growth by manual measurement of green leaf area (LA) every 5 days, starting at 10 days after emergence (DAE). Plants in Full conditions showed an increase in the rate of leaf initiation compared with reduced nutrient treatments (Fig. 1B; S1; MZ66_Growth_Analysis in Supplemental File 1). At 25 DAE, Full and LowP plants had initiated ∼1 more leaf than LowN and LowNP plants (KW adj. *p* = 0.003. Dunn test at α = 0.05. Leaf number - Full: 5.6 ±0.2a; LowP: 5.0 ±0.22a; LowN: 4.0 ±0.22b; LowP: 4.1 ±0.2b. Here and below, we give model coefficients, standard errors and means groups assigned by Dunn test or Tukey as indicated). By harvest, plants in Full had ∼1.5 more leaves on than those in the stress treatments, with the stress treatments indistinguishable among themselves (KW adj. *p* = 0.09. Dunn test at α = 0.05. Leaf number 40 DAE - Full: 8.6 ±0.28a; LowP: 7.3 ±0.32b; LowN: 7.0 ±0.33b; LowNP: 7.0 ±0.29b). The first two leaves were fully expanded in all treatments when we started to collect measurements at 10 DAE, and they began to senesce early in the course of the experiment, reflected by a loss of LA (Fig. 1C,D). Second leaves showed equivalent LA in all treatments (Leaf 2 LA - KW adj. *p* > 0.05 for treatment at all time points) and began to senesce at the same time (∼30 DAE; Fig. 1D). Senescence began earlier in first leaves than second leaves (∼20 DAE), and was more rapid under LowN and LowNP than in the other conditions (Fig. 1C. Leaf 1 - LA KW adj. *p* < 0.001 at 25 and 30 DAE. Dunn test at α = 0.05. First leaves senesced completely by day 25 under LowN and LowNP, but not until 40 DAE under Full and LowP). Third leaves were present at the first time point, continuing to grow until ∼20 DAE, with no difference in LA between treatments (Fig. 1E. Leaf 3 LA - KW adj. *p* > 0.05 for treatment at all time points). Later leaves were initiated during the experiment, showing growth differences between treatment (Fig. 1F-J; S1; MZ66_Growth_Analysis in Supplemental File 1). Treatment differences became more dramatic with each leaf to be initiated. In fourth leaves, we observed a mild treatment effect from ∼10 days after leaf expansion (Leaf 4 LA 20 DAE - KW adj. *p* = 0.044), leaves of the LowNP plants having a lower surface area than those of the other treatments (Fig. 1G; Dunn test at α = 0.05). Differences in the later leaves were evident within 5 days after initiation, the timing of initiation also becoming delayed in the low nutrient treatments. By the sixth and seventh leaves, we observed a difference between Full, LowP and LowN/LowNP treatments (Fig. 1H,I; S1; MZ66_Growth_Analysis in Supplemental File 1; leaf 6 LA, 30 DAE - KW adj. *p* < 0.001; leaf 7 LA, 35 DAE - KW adj. *p* < 0.001; Dunn test at α = 0.05; differences maintained until 40 DAE). Plants in the Full treatment also produced eighth (Fig. 1J) and some ninth (not shown) leaves. In addition to photosynthetic surface area, plant stature clearly differed among treatments, as captured by measurement of stem height (Fig. 1K; S1; MZ66_Growth_Analysis in Supplemental File 1). By 20 DAE, Full and LowP plants were taller than LowN and LowNP plants (Fig. 1K. Stem height - KW adj. *p* = 0.011; Dunn test at α = 0.05. Stem height 20 DAE - Full: 5.76 cm ±0.48a; LowP: 5.59 cm ±0.53a; LowN: 4.09 cm ±0.54b; LowNP: 3.83 cm ±0.48b), a pattern maintained until harvest. Treatment had a significant effect on total leaf area by 20 DAE (Fig. 1L; Square root transformed total LA 20 DAE - KW adj. p < 0.05; Dunn test at α = 0.05. Full: 7.16 cm ±0.46a; LowP: 6.72 cm ±0.52a; LowN: 5.82 cm ±0.53ab; LowNP: 4.72 cm ±0.47b). By harvest, the four treatments could be distinguished by total leaf area (Fig. 1L; Square root transformed total LA 40 DAE - KW adj. p < 0.001; Dunn test at α = 0.05. Full: 19.90 cm ±1.02a; LowP: 13.55 cm ±1.14ab; LowN: 9.15 cm ±1.17bc; LowNP: 8.35 cm ±1.04c). We used the slope of a linear fit through the plot of square root transformed total LA on time as an estimate of growth in each treatment and observed differences between all four treatments, with a ranking of Full, LowP, LowN, LowNP (Fig. S2; MZ66_Endpoint_Analysis in Supplemental File 1. Growth - KW adj. p < 0.001; Dunn test at α = 0.05. Full: 0.45 cm/day ±0.02a; LowP: 0.34 cm/day ±0.02ab; LowN: 0.25 cm/day ±0.02bc; LowNP: 0.22 cm/day ±0.02c)

**Figure 1.**
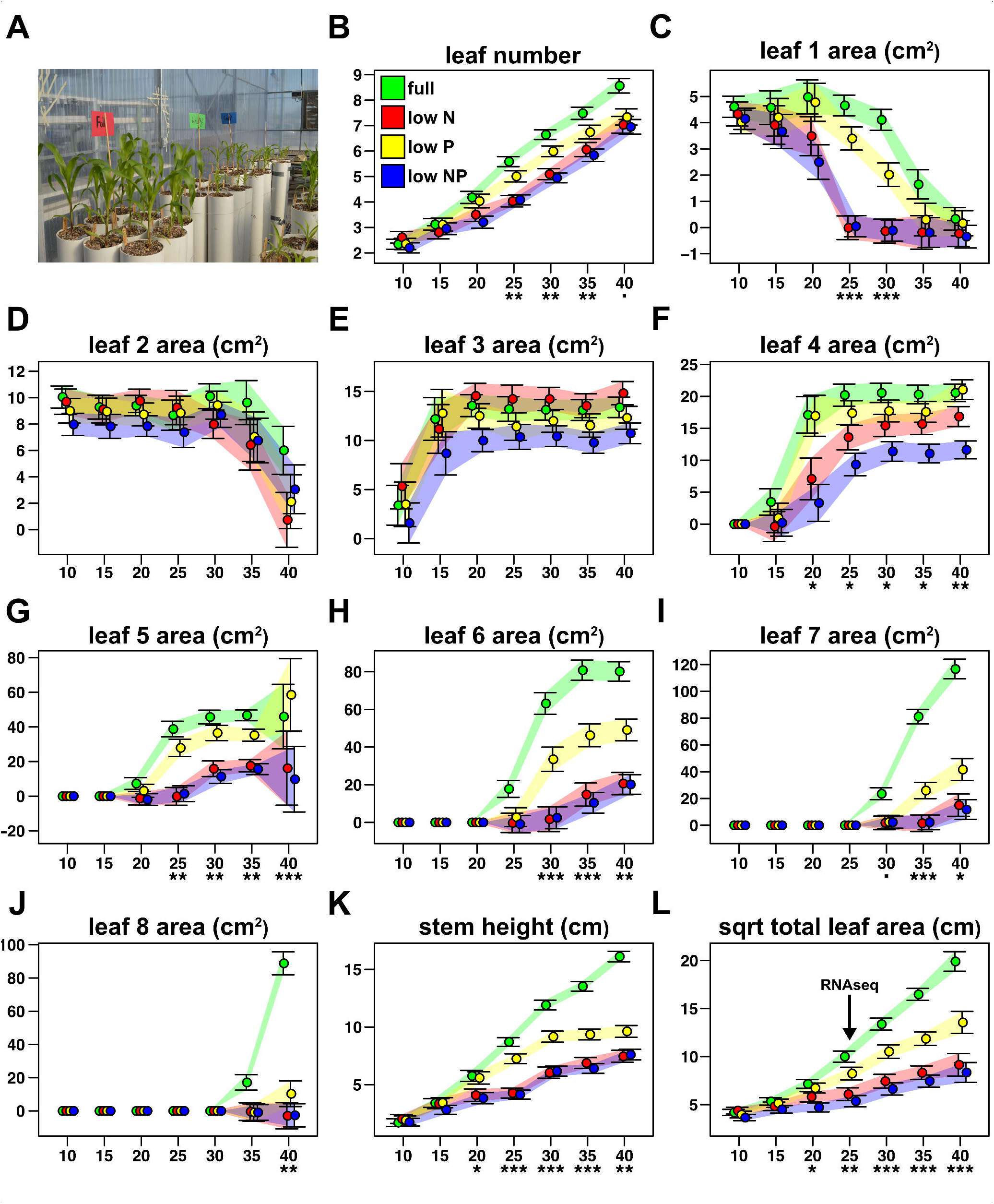
Total leaf area is reduced under low N and P availability from day 25 after emergence. A) General view of plant growth system. B) The number of fully expanded green leaves in plants grown in Full, LowP, LowN or LowNP treatments. Data collected every 5 days from 10 days-after-emergence (DAE) until day 40. Points show the coefficient estimated for each treatment, with bars extending +/- 1 standard error (SE). Colored polygons follow SE bars. Significance of treatment effects on a given day (Kruskal-Wallis test; p-value adjusted for multiple tests) are indicated below the x-axis as *** p <0.001, ** p <0.01, * p <0.05,. p<0.1. C-L) Coefficients of fully expanded green leaf area and stem height as B for further non-destructive traits. The arrow in L indicates the point at 25 DAE selected for the harvest of plants grown in the subsequent RNA sequencing experiment.

At 40 DAE, plants were harvested by careful removal from the PVC tubes and endpoint measurements taken for roots and the aerial portions of the plant. Plants under Full treatment were clearly larger than those under the stress treatments, while the three stress treatments were partially separated (Fig. 2A, B. MZ66_Endpoint_Analysis in Supplemental File 1. Root fresh weight - KW adj. *p* < 0.001; Dunn test at α = 0.05. Full: 18.07 g ±1.36a; LowP: 10.13 g ±1.52ab; LowN: 5.17 g ±1.55bc; LowNP: 3.62 g ±1.37c. Shoot fresh weight - KW adj. *p* < 0.001; Dunn test at α = 0.05. Full: 18.55 g ±1.00a; LowP: 6.14 g ±1.13ab; LowN: 2.48 g ±1.15bc; LowNP: 1.91 g ±1.02c). The root:shoot ratio (based on fresh weight) was greater under stress treatments, with an increase of 2.2-, 1.7- and 1.9-fold with respect to Full under LowN, LowP and LowNP, respectively (Fig. 2C). Plants under LowP and LowNP showed greater root system depth (RSD) than those under Full or LowN conditions, although the treatment effect was not significant (RSD KW adj. *p* = 1). Specific root depth (SRD), calculated as RSD over total root fresh weight, did vary significantly among treatments, with LowN and LowNP showing higher values than LowP and Full treatments (Fig. 2D. MZ66_Endpoint_Analysis in Supplemental File 1. SRD - KW adj. *p* < 0.001; Dunn test at α = 0.05. LowNP: 19.12 cm/g ±1.46c; LowN: 13.37 cm/g ±1.65bc; LowP: 9.49 cm/g ±1.62ab; Full: 5.02 cm/g ±1.44a).

**Figure 2.**
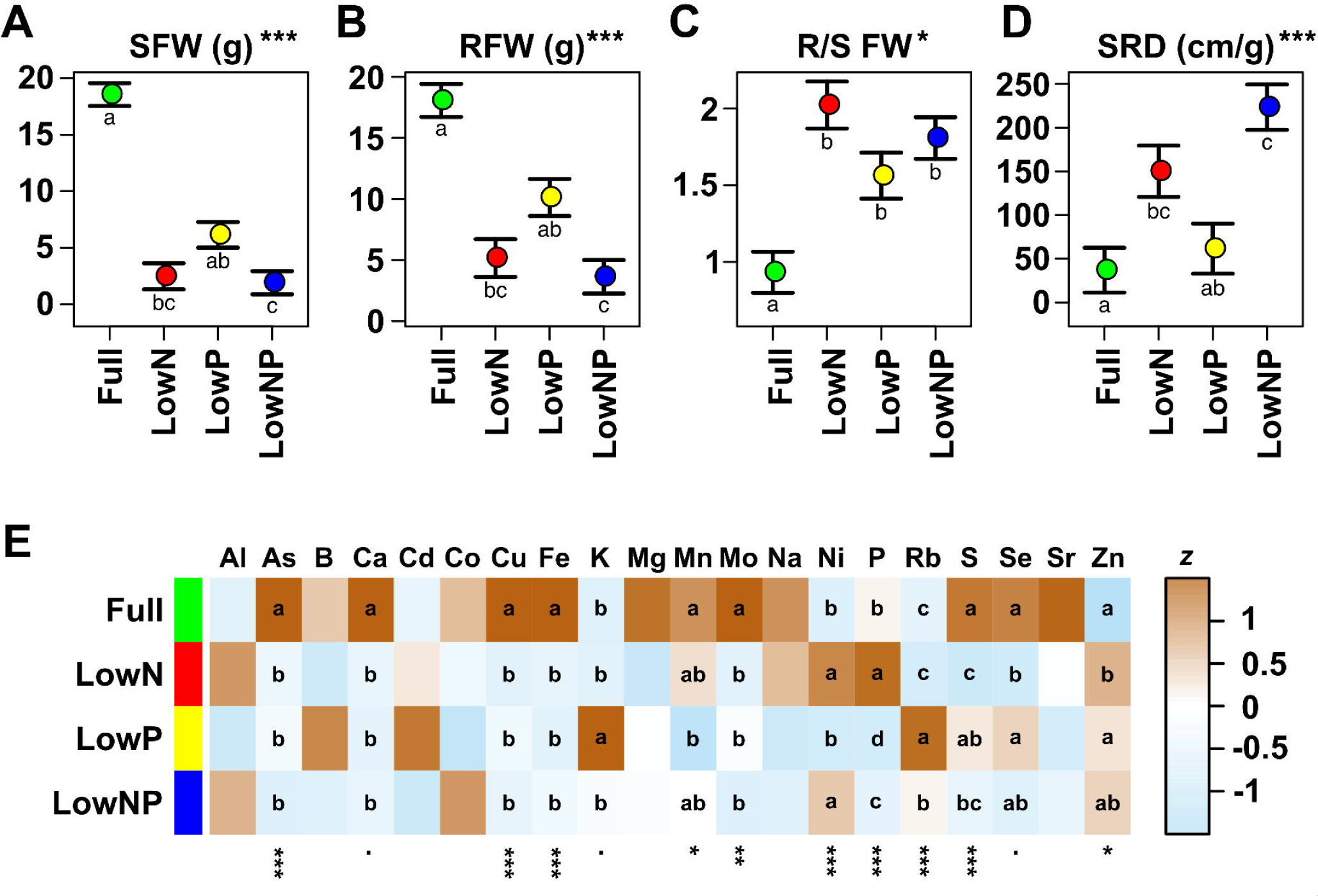
Low N and P availability alters the relative growth and element profile. A) Root fresh weight at harvest (RFW, g; estimated coefficient and associated standard error) of plants grown in Full, LowN, LowP or LowNP. The significance of the treatment effect is shown as *** p <0.001, ** p <0.01, * p <0.05, p<0.1 (Kruskal-Wallis test; p-value adjusted for multiple tests). Lowercase letters indicate significant (p<0.05) pairwise differences (Dunn test). B-D) As A, showing shoot fresh weight (SFW, g), the ratio of RFW/SFW (RS) and specific root depth (SRD cm/g), respectively. E) Heat map representation of total ion concentration for 20 named elements (z, concentration standardized within row). The significance of the treatment effect on concentration is shown as *** p <0.001, ** p <0.01, * p <0.05, . p<0.1 (ANOVA; p-value adjusted for multiple tests). Lowercase letters indicate significant (p<0.05) pairwise differences (Tukey).

To further characterize differences in root system architecture (RSA) among nutrient treatments, we photographed the roots of each plant and processed the images using GiaRoots analysis software (Galkovskyi et al. 2012) to extract a series of root features. Nutrient treatment had a significant effect on several root features related to overall root system size (Fig. S3; MZ66_Giaroots_Analysis in Supplemental File 1) including network area, perimeter and volume and the maximum and median number of roots crossing a horizontal line in a vertical scan (see [55] for a complete description of root features). The Full treatment was associated with the largest, most solid root systems, followed by LowP, LowN and LowNP. We also saw a significant treatment effect on the ratio of the minor/major axes (EAR) of an ellipse fitted around the root system (EAR - KW adj. *p* = 0.016; Dunn test at α = 0.05. Full: 0.46 ±0.03a; LowP: 0.40 ±0.04a; LowN: 0.36 ±0.04a; LowNP: 0.29 ±0.03b). EAR reflects the tendency to relatively narrower but deeper root systems in the low nutrient treatments. In comparison to leaf growth and aerial biomass traits, root features were more similar in the single LowN and LowP treatments, and there was greater evidence of a partially additive effect in the combined LowNP treatment (Fig. S3; MZ66_Giaroots_Analysis in Supplemental File 1. *e.g.* network volume - KW adj.*p* = 0.009; Dunn test at α = 0.05. Full: 19.40 ±1.68a; LowP: 12.43 ±1.89ab; LowN: 10.10 ±1.92bc; LowNP: 7.23 ±1.71c).

### The leaf ionome was modified under low N and P treatments

To further characterize the impact of N and P limitation on early growth in maize, we quantified the total concentration of twenty different elements in the leaf tissue using inductively coupled plasma mass spectrometry (ICP-MS). The technique used did not determine N concentration. We detected a significant (ANOVA adj. p < 0.05) effect of treatment on the concentration of ten of the elements quantified (Fig. 2E, S4; MZ66_Ionomics_Analysis in Supplemental File 1). We observed both decreases and increases in concentration for different elements, indicating that the effects could not be explained solely based on changes in root:shoot ratio. In line with previous studies [15, 22, 23, 38], we observed an increase (1.8 fold) in leaf total P concentration of plants grown under LowN compared with Full (P concentration - ANOVA adj.*p* < 0.001; Tukey test at α = 0.05. LowN: 3543 ppm ±110a; Full: 1983 ppm ±96b; LowNP: 1174 ppm ±98c; LowP: 734 ppm ±108d). Unsurprisingly, total P concentration was lower under LowP (734 ppm). More remarkably, total P concentration was higher under LowNP (1174 ppm) than in LowP, although LowNP plants were also smaller than those under LowP. We also saw a significant increase in Ni concentration under LowN and increases in K and Rb concentration under LowP (Fig. 2E, S4; MZ66_Ionomics_Analysis in Supplemental File 1).

### The transcriptional response to P starvation is repressed under N limitation

Based on our initial characterization, we selected 25 DAE - the point at which we first saw a significant treatment effect on growth across leaves (Fig. 1) - for transcriptional profiling. We grew a second set of plants under the same nutrient conditions as used previously, harvesting total roots and pooled leaf blades at 25 DAE from two individuals per treatment for RNA extraction and sequencing. Sequencing reads were aligned to the maize (var. B73, ref-gen V3) transcript set and collapsed at the gene level to obtain read count data. We analyzed count data from all treatments and both tissue in a single linear model to identify significant effects of LowN, LowP or their interaction on gene expression. A total of 1,555 genes were identified to be N/P regulated (false discovery rate [FDR] all nutrient terms < 0.01; |log_2_ fold change [LFC]| > 1 for at least one nutrient-associated model term; MZ67_DEG_set in Supplemental File 2). Regulated genes were further classified as upregulated or downregulated in different tissue/treatment combinations by the sign and magnitude (|LFC| > 1) of pairwise differences with respect to the Full treatment in the relevant tissue (MZ67_DEG_set in Supplemental File 2).

A similar number of genes were upregulated as were downregulated; a greater number of regulated genes were detected in leaves than roots (Fig. 3A-D). We compared the transcriptional response to the treatments by tissue and sign of the effect (up or down). There was little overlap between the responses to LowN and LowP single stress treatments (Fig. 3A-D. *e.g.* of a combined total of 737 genes upregulated in the leaf between LowN and LowP, only 30 were shared). When presented with the combined LowNP treatment, plants broadly followed the LowN response pattern: most regulated genes were shared between LowN and lowNP; very few genes that were regulated under LowP showed similar regulation under lowNP (Fig. 3A-D). This trend was evident in both leaves and roots, and among both up- and down-regulated genes.

**Figure 3.**
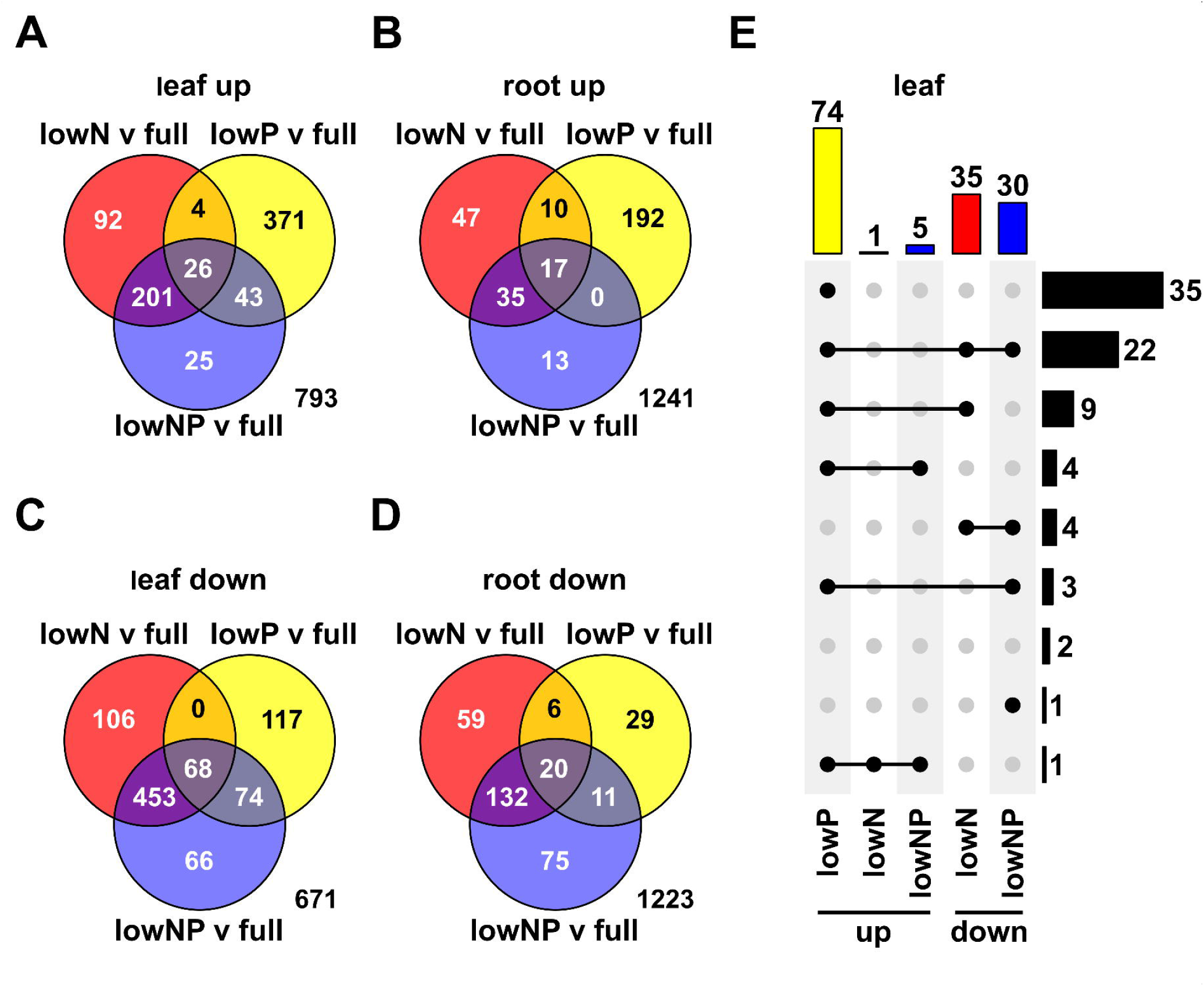
Transcriptional responses to LowN and LowP are distinct. A) Grouping of 1,555 N/P regulated genes (MZ67 DEG set in Supplemental File 2) with respect to upregulation in leaves under LowN (red), LowP (yellow) or combined LowNP (blue) in comparison with Full. B - D) As A, with respect to downregulation in leaves, upregulation in roots and downregulation in roots, respectively. E) Upset diagram classifying the 81genes that showed significant NxP interaction. Filled circles connected by line segments indicate common set membership. Colored bars and numbers at the top of the plot show the size of each set, colored by treatment comparison (colors as A-D). Black bars on the right of the plot indicate the number of genes in each intersection.

Our model included an NxP interaction term. Although our power to detect interaction effects was no doubt limited by the level of replication, we were able to identify 81 NxP interaction genes (FDR all NxP terms < 0.05; |LFC| > 1 for at least one nutrient interaction model term. MZ67_NxP_set in Supplemental File 2); *i.e.* genes regulated by the availability of one nutrient in a manner conditional on the availability of the second. We explored the distribution of these 81 genes across the sets of upregulated and downregulated genes (pairwise |LFC| > 1) from the different treatments and tissues (Fig. 3E). The majority (74 of 81) of NxP genes were upregulated in the LowP single treatment in leaves (Fig. 3E; MZ67_NxP_set in Supplemental File 2). Only 5 of these 74 leaf LowP induced NxP genes were also upregulated under LowNP. Furthermore, 35 and 25 of these 74 were downregulated in the leaves under LowN and LowNP, respectively (Fig. 3E; MZ67_NxP_set in Supplemental File 2). A similar pattern was seen in roots - 60 of the 81 NxP genes were upregulated in roots under LowP, 55 in common with the leaves; none of the 81 NxP genes were upregulated in the roots under lowN or lowNP; ten and 9 root LowP induced NxP genes were downregulated in roots under LowN and LowNP, respectively (MZ67_NxP_set in Supplemental File 2). These observations suggested that the responses to LowP and LowN were not only distinct, but antagonistic, and that under LowNP the pattern seen under LowN dominated. Although NxP interaction were not supported statistically beyond these 81 genes, a similar global pattern was seen across the complete 1,555 gene set. Of 444 genes up-regulated in leaves under LowP (pairwise LFC > 1), only 30 were up-regulated under LowN, while 178 were down-regulated (Fig. 4A; MZ67_DEG_set in Supplemental File 2). Similarly, of these 444, only 69 were up-regulated under combined LowNP, with 121 down-regulated (Fig. 4A, S5A-B. MZ67_DEG_set in Supplemental File 2).

**Figure 4.**
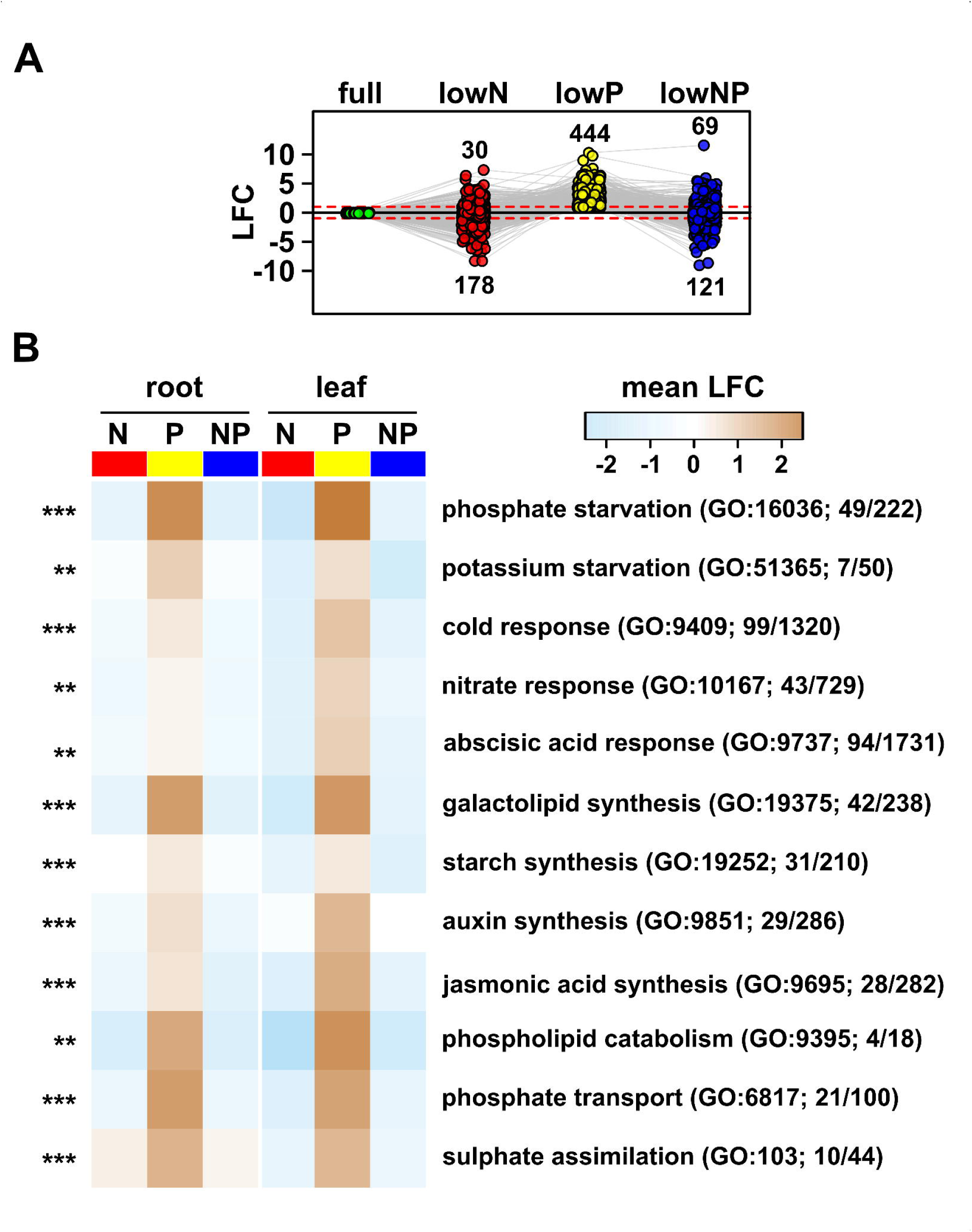
Transcript induction under LowP is repressed by LowN. A) Reaction norm plot of differential transcript accumulation (log_2_ fold change, LFC) for 444 genes induced by lowP in the leaf with respect to Full (green) under LowN (red), LowP (yellow) and combined LowNP (blue). Numbers indicate the count of genes above/below the +/- 1 LFC threshold. B) Heat map showing the mean LFC in each treatment of genes associated with selected GO terms enriched in the 1,555 gene set (Supplemental File 2). LFC calculated with respect to Full, separately for roots and leaves. GO term names are abbreviated. GO term identifiers are given in parentheses along with the number of genes assigned in the test set over the total number in the GO term. The significance of GO term enrichment is indicated to the left of the heat map as *** p <0.001, ** p <0.01, * p <0.05.

To gain insight into the functional consequences of the transcriptional responses, we examined “classical genes” (a curated set of ∼5,000 well-annotated genes, many linked with existing functional data: maizegdb.org/gene_center/gene) in our regulated gene set. We supplemented the classical set with a number of additional annotations [56, 57] based on identification of maize orthologs of high-interest candidate genes, notably members of the maize SPX-domain and PAP gene families (Fig. S5C; [58]). The SPX-domain family proteins have been clearly linked with crosstalk in N-P signaling in *Arabidopsis* and rice [42, 44, 50], but the family has not been previously annotated in maize. We therefore identified the complete set of SPX-domain protein encoding genes from maize and assigned a nomenclature based on phylogenic analysis that we use below (Fig. S6). The behavior of the top thirty (ranked by FDR) regulated classical genes mirrored the global trend - namely, strong induction under LowP that was absent, or shifted to repression, in LowN or LowNP (Fig. S5C). The top classical genes encoded functions previously associated with the PSR [59–62], including PHT1 high-affinity phosphate transporters, PAPs, lipid-remodeling enzymes and members of the SPX domain family (Fig. S5C, S6). We further examined functional patterns using Gene Ontology (GO) analysis of the complete 1555 regulated gene set (MZ67_GO_Analysis in Supplemental File 2). We calculated the mean LFC with respect to full nutrients of the regulated genes belonging to each enriched GO set under the three nutrient treatments, in roots and leaves (Fig. 4B; MZ67_GO_Analysis in Supplemental File 2) As for the single gene analysis, we observed a signature of upregulation under LowP associated with downregulation under LowN. Of the top 50 GO sets by *p* value for enrichment, the mean LFC under LowP was positive for 46 sets in roots and for all 50 in the leaves; for the same sets the mean LFC was negative in all but two cases in LowN and LowP treatments, in both root and leaves (MZ67_GO_Analysis in Supplemental File 2). This pattern extended from the general *cellular response to phosphate starvation* term (GO: 16036) to specific processes such as *galactolipid biosynthetic process* (involved in lipid-remodeling under P starvation. GO: 19375) and synthesis of the hormones auxin (GO: 9851) and jasmonic acid (GO: 9695; Fig. 4B).

### Mild N stress is sufficient to repress the P starvation response

Although LowN and LowP treatments were adjusted to 9 and 3% of the Full concentration, respectively, it was evident by 40 DAE that the LowN treatment produced a greater limitation on growth than LowP. As such, we speculated that the dominance of the LowN transcriptional response under the combined NP treatment was simply a consequence of the greater severity of the LowN stress. To address this hypothesis, we grew an additional set of plants under high and low P (P5 and P1, respectively; our original Full and LowP levels) in combination with five different levels of N (N5 to N1, high to low; the extremes corresponding to our previous Full nutrient and LowN treatments) to measure the accumulation of selected transcripts. As for our whole transcriptome experiment, we harvested plants at 25 DAE (Fig. 1). We measured shoot and root fresh weight and again saw that the single stress combination N1P5 reduced growth more than the complementary N5P1 treatment (Fig. 5A, B). At intermediate N availability, however, we could observe different combinations of N and P with equivalent growth: *e.g.*, N4P5 was indistinguishable from N5P1 in terms of shoot fresh weight. To evaluate the impact of N availability on the PSR, we used real-time PCR to quantify the expression of a panel of selected genes. We first assayed the well-characterized N responsive genes *Nir-a* (GRMZM2G079381) and *Npf6.6* (GRMZM2G161459), encoding a nitrate reductase and a nitrate/peptide transporter [17, 51], respectively, to confirm the impact of the N treatments. As previously shown and as observed in our transcriptome data (MZ67_DEG_set in Supplemental File S2), *Nir-a* and *Npf6.6* were down-regulated in reduced N treatments (*Nir-a* is expressed predominantly in leaf tissue. Fig. 5C, D; MZ95_DE_analysis in Supplemental File S3). The accumulation of *Nir-a* and *Npf6.6* decreased from N5 to N1 treatments, indicating a progressive impact on plant N status and signaling (Fig. 5C, D). Interestingly, expression of *Npf6.6* was also induced in the roots under P1, this response being most pronounced at N5. We then assayed four PSR genes, selected based on previous reports and our transcriptome data: *Pht1;9*, *Pht1:13* phosphate transporter genes in roots [27], the *Mfs2 SPX-*family gene in leaves, and the *Pap10* purple acid phosphatase gene in both roots and leaves [58] All four PSR genes were strongly induced by P1 under N5 conditions (Fig. 5C-D; *Mfs2* 1.8-fold increase N5P1/N5P5 in leaves; *Pap10* 1.85-fold increase N5P1/N5P5 in leaves, 4.93-fold increase in roots; *Pht1;9* 4.33-fold increase N5P1/N5P5 in roots; *Pht1:13* 4.82-fold increase N5P1/N5P5 in roots). However, once N availability was reduced to N4, the level of PSR transcript accumulation under P1 was reduced (Fig. 5C, D; MZ95_DE_analysis in Supplemental File S3). At N3 and below, P1 induction of PSR genes was absent. Interestingly, *Mfs2* and *Pap10* showed a level of constitutive expression in leaves under N5P5 conditions that was reduced by N limitation (*Mfs2* 2.10- and 14.72-fold reduction in N4P5 and N1P5, respectively; *Pap10* 4.94- and 19.70-fold reduction in N4P5 and N1P5, respectively; Fig. 5C-D; MZ95_DE_analysis in Supplemental File S3).

**Figure 5.**
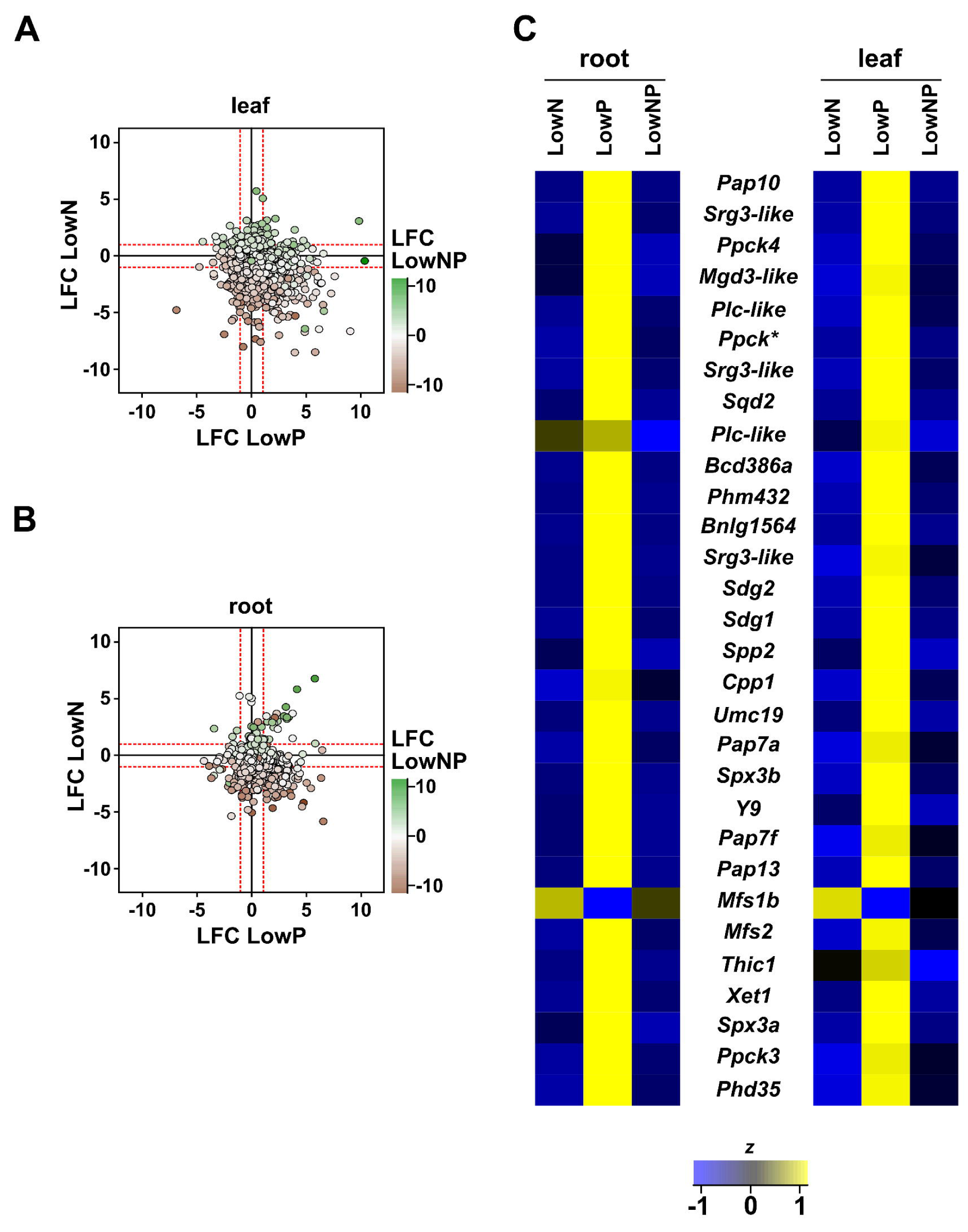
Moderate N stress is sufficient to repress the low P response. A) Representative 25-day-old maize seedlings grown across five levels of N availability (N5 to N1, high to low) and two levels of P availability (P5, high and P1, low). B) Shoot fresh weight of maize seedlings grown as A. Boxes show 1st quartile, median and 3rd quartile of 4 biological replicates. Whiskers extend to the most extreme points within 1.5x box length; outlying values beyond this range are not shown. Letters indicate groups based on HSD Tukey (p < 0.05). Transcript accumulation (relative abundance) determined by real-time PCR for C) leaves and D) roots of 25-day-old maize seedlings grown as A. Median of 5 biological replicates. *Pap10-Purple acid phosphatase10*, GRMZM2G093101; *Pht1;9* - *Phosphorus transporter1;9*, GRMZM2G154090; *Pht1;13 - Phosphorus transporter1;13,* GRMZM2G070087; *Mfs2* - *ZmSPX-MFS2*, GRMZM2G166976; *Npf6.6* - *Nitrate/peptide Transporter6.6*, GRMZM2G161459); *Nir-a* - *nitrite reductase-a*, GRMZM2G079381).

### P concentration in the leaves responds to both P and N availability in the substrate

Previous studies and our observations at 40 DAE showed an increase in total P concentration in the leaves of young plants grown under N limitation [15, 22, 23, 38]. As such, the antagonism observed between transcriptional responses to our LowN and LowP treatments might be driven by downregulation of PSR genes in response to higher cellular P concentration. To investigate this possibility, we quantified total P concentration using ICP-MS in the roots and leaves of the plants in our N-dose experiment (MZ95_Ion_Concentration_Analysis in Supplemental Table S3). We again observed an increase in total P concentration in both leaves and roots as N was reduced, in either P1 or P5 (Fig. 6A). However, the increase over the N5-N3 range was minimal (P concentration root. Tukey test at α = 0.05. N5P5: 1035 ppm ±100ab; N3P5: 1082 ppm ±82ab; N5P1: 809 ppm ±32b; N3P1: 1139 ppm ±55ab; P concentration leaf. Tukey test at α = 0.05. N5P5: 2525 ppm ±103efg; N3P: 3545 ppm ±203abc; N5P1: 2064 ppm ±81g; N3P1: 2277 ppm ±86fg), suggesting that total P concentration cannot explain the strong effects on gene expression we saw over the same range.

**Figure 6.**
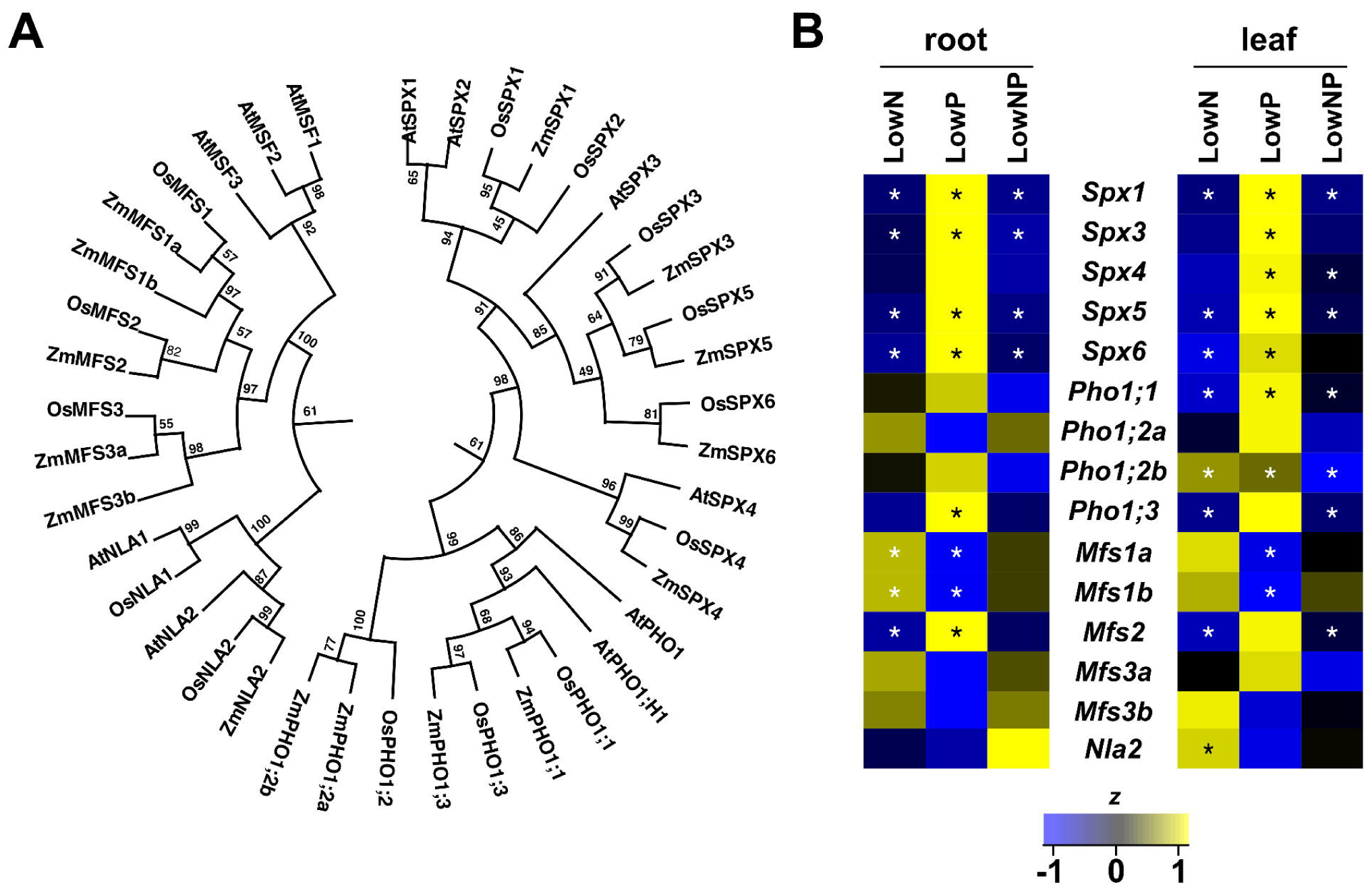
P accumulation responds to P and N availability. A) root and B) leaf P concentration (ppm dry mass) of 25-day-old maize seedlings grown across five levels of N availability (N5 to N1, high to low) and high (green points and trace) and low (yellow points and trace) levels of P availability. Large points show treatment medians; small points show individual (4) biological replicates. Dashed lines show best fit from a multiple regression model. Asterisks represent statistical significance of model terms (p value ≤ 0.001 ***; 0.001-0.01 **; 0.01-0.05 *). N, P main effect of N and P, respectively. NP, NxP interaction term. Lowercase letters indicate significant (p<0.05) pairwise differences (Tukey).

## DISCUSSION

To explore the interaction between N and P signaling pathways in maize, we characterized transcriptional responses in roots and leaves to low N, low P and combined low NP stress. We observed responses to our LowN and LowP treatments to be distinct and antagonistic. Furthermore, under combined LowNP, the LowN pattern dominated and the classic PSR was absent, even though plant growth was partially P limited (as determined by phenotypic comparison to plants grown under the single LowN stress). Although there were differences at the level of individual genes, our LowN and LowP single stress results are in broad agreement with a previous report in which a similar antagonism was observed, and many classic PSR genes were seen to be down-regulated under LowN [15]. The rationale, or potential adaptive value, of such antagonism is not clear.

N is typically found deeper in the soil than P, reflecting differences in mobility. Consequently, a root system optimized to access P in the topsoil will be less suited to N acquisition, and *vice versa* [65–67]. In addition, the optimal pattern of root branching and root length is different for acquisition of N or P [65–67]. We did not detect dramatic differences in RSA between LowN and LowP treatments at 40 DAE, although the growth system, the relatively young age of the plants, and the severity of stress may have limited the expression of potential root developmental responses. Nonetheless, the antagonistic regulation of genes associated with hormone signaling (*e.g.*, genes belonging to GO terms GO:9851 *auxin biosynthetic process*, GO:9695 *jasmonic acid biosynthetic process*; GO:9735 *response to cytokinin*) may mirror the differing demands placed on plant architecture by N and P limitation.

Once acquired, the efficiency of internal P use can be maximized by remobilization to the part of the plant where need is greatest over the growing season [3, 4]. PAPs remobilize P by releasing inorganic P from organic compounds. Induction of PAP encoding genes and increased PAP activity is a classic component of the PSR across the tree of life, including *Arabidopsis* [68, 69], rice [70] and maize [58]. We observed several *Pap* genes to be upregulated under LowP in both roots and leaves. In addition to remobilizing P within the plant, PAPs are also secreted to the rhizosphere, enhancing the availability of inorganic P for uptake [68, 69, 71]. *Pap10* was one of the most strongly regulated genes in our analysis. Reflecting the global pattern, *Pap10* was strongly induced by LowP, but only in N4-N5 conditions. Furthermore, *Pap10* showed a constitutive level of expression in our Full nutrient condition that was reduced by lowering N availability. Genes linked to lipid remodeling - the replacement of membrane phospholipids by galactolipids or sulfolipids under P starvation [4, 72, 73] - followed a similar trend. Downregulation of constitutively expressed PSR-associated genes by single low N treatments has been previously reported in four commercial maize hybrids and two maize inbred lines [28, 32, 74]. One study that did not report such downregulation of PSR gene also found no evidence of the downregulation of N assimilation genes typically associated with N starvation, indicating that the precise nature and timing of the treatment are important [75]. A similar down-regulation of PSR genes occurs in rice under prolonged N starvation [50], but not within the first 12 hours of shift to N starvation conditions [5], although a metabolic response can occur as early as 1 hour after such a shift [76]. Our observations that the negative impact of low N availability on PSR gene expression dominates in the combined LowNP treatment implies that, under this dual stress, maize plants are failing to activate well-defined aspects of the PSR, such as P remobilization or lipid remodeling. In the future, it will be informative to assay PAP activity and lipid composition at low N and low P availability.

Our study confirmed previous observations of P hyper-accumulation in maize leaves under N limitation [15, 28], an effect also reported in rice and *Arabidopsis* [42, 77]. Initially, we considered the hypothesis that down-regulation of PSR genes in LowN was a secondary response to an increase in total internal P concentration. However, LowNP conditions downregulated PSR genes even when low P availability prevented accumulation of total P to the concentration seen under LowN conditions. Significantly, mild N limitation (N4) was sufficient to suppress induction of PSR genes under LowP with no change in internal total P concentrations. Plants perceived N reduction from N4 and below, as demonstrated by the reduced accumulation of *Nir-a* transcripts, a well characterized marker of plant N status [78]. Overall, our data support an N-mediated impact on PSR via modified signaling or P partitioning, rather than as the secondary effects of total internal P hyper-accumulation.

Currently, it is difficult to reconcile PSR repression and P hyper-accumulation. It would be informative to examine earlier stages of plant growth for evidence of a transient induction of PHT1 transporter encoding genes under LowN, although no such signal has been previously reported in comparable experiments in maize or other plants, nor in experiments using a transfer from replete to N starvation conditions [76, 79]. PHT1 transporters are subject to regulation at the post-translation level [45, 80, 81] and measurement of protein levels and localization would provide a fuller picture, as would quantification of root P permeability and P uptake. In rice, it has been reported that the roots of plants grown under N starvation show increased permeability to inorganic P [38]. The balance between P concentration in the leaves and P uptake by the roots is maintained by systemic signaling through the mobile microRNA miR399 [82, 83]. As P becomes limiting in the shoots, miR399 is produced and travels to the roots to target transcripts encoding the PHOSPHATE2 (PHO2) E2 ubiquitin conjugase, in turn promoting accumulation of PHT1 transporters [84–86]. Previous reports have shown that miR399 expression in maize can increase in N starvation, although the effect depends on both the nature of the N treatment and the length of exposure [87, 88].

Study of NP crosstalk in *Arabidopsis* and rice has highlighted the importance of the SPX protein family. Although first described as regulators of P homeostasis [92], SPX and SPX-RING proteins have subsequently been linked with N signaling [42, 44, 50]. We identified 15 SPX-domain family genes in maize, the same as in rice, grouped into the four previously reported classes (*SPX*, *SPX-EXS*, *SPX-MFS* and *SPX-RING*; [64]). N and P availability regulated transcript levels across the SPX family, consistent with a role in the integration of N and P signaling pathways (Fig. S6). Transcripts encoding members of the single SPX domain class responded positively to LowP in both roots and leaves, as has been seen previously in *Arabidopsis* and rice [94, 95]. In rice, over-expression of *OsSPX1* and *OsSPX6* suppresses the PSR, suggesting that they may act in a negative-feedback loop. Conversely, under-expression of *OsSPX1* and *OsSPX6* leads to increased P accumulation through upregulation of genes involved in P uptake [95, 96]. The rice SPX4 protein exerts a further negative control on the PSR by sequestering the MYB transcription factor PHR2 in the cytosol, preventing its translocation into the nucleus and activation of target genes [49]. Under P starvation, SPX4 is degraded, freeing PHR2 to activate the PSR. It has recently been reported that SPX4 turnover in rice requires the activity of the NRT1.1b [50]. Given that the abundance of NRT1.1b itself is N responsive, the NRT1-SPX4 module represents a point of integration between N and P signaling pathways.

Hyperaccumulation of P under N limitation indicates an uncoupling of P uptake from leaf P concentration [84–86]. Similar uncoupling occurs in *Arabidopsis* mutants underexpressing the SPX-EXS gene *PHO1*, in parallel with changes in subcellular partitioning of P between vacuolar stores and the cytosol [97]. The maize genome encodes two co-orthologs of the *Arabidopsis PHO1* - *Pho1;2a* and *Pho1;2b* [93]. We found both *Pho1;2a* and *Pho1;2b* to show evidence of downregulation under LowN, potentially contributing to changes in P partitioning. While our observations suggest that changes in total internal P concentration cannot explain the observed effect of N limitation on the PSR, we do not have data on the level of P in the cytosol itself. A second group of SPX proteins, the SPX-MFS proteins, plays a more direct role in regulating cytosolic P concentration by mediating P influx into the vacuole [98, 99]. Under P starvation, *OsSPX-MFS1* and *OsSPS-MFS3* are down regulated, consistent with retaining more of the total internal P pool in the cytosol for direct use [87]. In contrast, *OsSPX-MFS2* is upregulated under P starvation, and may be acting differently [100, 101]. The MFS2 protein was not identified in a screen for vacuolar P efflux transporters [99], suggesting that it is not simply working antagonistically to MFS1 and MFS3. In maize, we found both *Mfs1* and *Mfs3* to be encoded by two genes, with both paralogs of each pair down regulated under LowP in the leaves, indicating a similar function to the rice genes. *Mfs2* was found to be a single copy gene in maize, and, as in rice, to be upregulated under LowP. It will clearly be greatly informative to functionally characterize the link between the maize SPX-domain proteins and N-P signaling.

## CONCLUSIONS

A reduction in N availability suppresses the PSR in young maize plants. Somewhat paradoxically, low N availability also results in an increase in internal P concentration – although not to levels that might explain the repression of low P responsive genes. In cultivated fields, P limitation may coincide with low N availability. As such, maize may grow without the classical low P response of model systems, making us rethink our current understanding of acclimation to P starvation. Further work is needed to evaluate the nature of the transcriptional PSR in maize under cultivation. We might also consider the merits of biotechnological manipulation to enhance low P responses under low N conditions.

## METHODS

### Plant material and growth conditions

Plants in this study were maize (*Zea mays* ssp. *mays* var. W22) wild-type segregants from a larger population segregating for the *Zmpho1;2-m1.1’* mutation, generated from the stock *bti31094::Ac* [93]. The original *bti31094::Ac* stock is available from the Maize Genetics Cooperation Stock Center. Genotypic analysis of the segregating population was as described previously [93]. Samples from individuals carrying the *Zmpho1;2-m1.1’* mutation were retained for future analysis. Plants were grown in the greenhouse in sand substrate with nutrient conditions maintained by a combination of fertilization with Hoagland solution [102]; 5mM KNO_3_, 0.25mM Ca(NO_3_)_2_, 2mM MgSO_4_, 1mM KH_2_PO_4_, 20μM FeC_6_H_6_O_7_, 9μM MnSO_4_, 1.2μM ZnSO_4_, 0.5μM CuSO_4_, 10μM Na_2_B_4_O_7_, 0.008μM (NH_4_)6Mo_7_O_24_), modified as described below and, were stated, by addition of 1.5% (v/v) of P-charged acidified alumina [103]. Hoagland N concentration was adjusted by substitution of KNO_3_ with KCl and CaCl_2_ [104, 105]. Hoagland solution was applied at 1/3 strength with the final N and P concentrations used in different experiments as stated below.

For growth to 40 days after emergence (DAE), 35 plants were evaluated in PVC tubes (15 cm diameter; 1m tall), planted in 4 groups, at intervals of one week. Tubes were filled with ∼17 litres of washed sand. In the upper third of the tube, soil was mixed with 1.5% solid-phase P buffer (alumina-P) [103] loaded with 209 μM KH_2_PO_4_ for Full treatments and 11 μM KH_2_PO_4_ for LowP treatments. Four imbibed seeds were planted at 4 cm depth per tube, thinned to a single plant a week after emergence. Plants were irrigated with distilled water up until 10 DAE after which Hoagland treatments were applied as a 1/3 strength solution, at a rate of 200 ml every third day, with final concentration: Full 1750 μM NO_3_; LowN 157.5 μM NO_3_; Full 333 μM KH_2_PO_4_; LowP 10 μM KH_2_PO_4_. During the growth period, plants were evaluated by non-destructive measurement of stem width, stem height, leaf number, and length and width of each fully expanded leaf. Stem height was measured from the soil to the last developed leaf collar. Measurements were collected every fifth day from 10 DAE. At 40 DAE, plants were removed from the tubes, minimizing damage to the root system, washed in distilled water and dried with paper towels before measuring root and shoot fresh weight. The cleaned root system was placed in a water-filled tub and photographed using a digital Nikon camera D3000. Raw images were individually processed using Adobe Photoshop CC (Version 14.0) to remove the background and obtain a good contrast between foreground and background non-root pixels. Processed images were scaled and analyzed using GiA Roots software [55]. Roots and shoots were placed in an oven at 42°C for a week before measuring dry weight and collecting samples for ionomic analysis (see below). The complete set of measurements collected is described in MZ66_Raw_Data in Supplemental File 1.

For growth up to 25 DAE, plants were grown in smaller PVC tubes (15 cm diameter, 50 cm tall). For the RNA-seq analysis, the top 30 cm of the 50 cm tube included 1.5% solid-phase P buffer (alumina-P [103]). The whole plant was harvested, separating the stem and leaves, a segment 2 cm above and below the crown roots and the remaining root system. Tissue was immediately frozen in liquid nitrogen and stored at −80 °C. Samples were homogenized with cooled pestle and mortar and aliquoted under liquid nitrogen for transcriptome analysis. For the N-dose experiment, plants grown in 50 cm tubes were irrigated with combinations of P at 10 or 333 μM (P1, P5; solid-phase P buffer was not used in this experiment), and N at 157.5, 233, 350, 875 or 1750 μM (N1 to N5). Leaf and root tissue were collected at 25 DAE for gene expression and ionomic analysis.

### Determination of elemental concentration by ICP-MS analysis

Ion concentration was determined as described previously by [106]. Briefly, root and shoot samples were analyzed by inductively coupled plasma mass spectrometry (ICP-MS) to determine the concentration of twenty metal ions. Weighed tissue samples were digested in 2.5mL concentrated nitric acid (AR Select Grade, VWR) with an added internal standard (20 ppb In, BDH Aristar Plus). Concentration of the elements Al, As, B, Ca, Cd, Co, Cu, Fe, K, Mg, Mn, Mo, Na, Ni, P, Rb, S, Se, Sr and Zn was measured using an Elan 6000 DRC-e mass spectrometer (Perkin-Elmer SCIEX) connected to a PFA microflow nebulizer (Elemental Scientific) and Apex HF desolvator (Elemental Scientific). A control solution was run every tenth sample to correct for machine drift both during a single run and between runs.

### Statistical analysis of plant growth and ionomic data

For plants grown to 40 DAE, traits were obtained from 34 individuals (one individual was removed as a clear outlier with poor growth). Individuals were distributed across nutrient treatments as: Full, n=7; LowN, n=5; LowP, n=9; LowNP, n=13, across 4 planting date. Traits included direct measurements and derived values (*e.g.*, total leaf surface area or biomass totals). Non-destructive measurements were repeated at 5-day intervals during the experiment. Destructive measurements were made for all 34 individuals at harvest. The data set include element concentrations determined by ICP-MS and root architectural traits extracted by image analysis, as described above. The dataset and analysis are presented in Supplemental File 1.

All statistical analysis was performed in R [107]. Full, LowN, LowP and LowNP were treated as four levels of a single treatment factor. For growth and endpoint data and GiaRoots features, we used R/stats::kruskal-test to assess the treatment effect on each trait with a non-parametric Kruskal-Wallis test. Element concentration was analyzed using ANOVA. In all cases, p-values were adjusted for multiple testing using the Bonferroni method with R/stats::p.adjust, applied separately to growth, endpoint, GiaRoots and element data sets. Where the treatment effect was significant (adjusted p < 0.05), we applied a pairwise *post hoc* test to identify differences between treatments: Dunn’s test (R/dunn.test::dunn.test [108]) for growth, endpoint and GiaRoots features and Tukey HSD for element data (R/agricolae::HSD.test [109]). For Dunn’s test results, letters were assigned to means groups using R/multcompView::multcompLetters [110]. For visualization, we used R/stats/lm to fit the model *trait value* ∼ 0 + *treatment* + *planting date* + *error*, extracting model coefficients and standard errors for plotting.

### RNA-sequencing analysis of differential gene expression

RNA-sequencing analysis was carried out on roots and leaves for the 4 nutrient treatments (Full, LowN, LowP and lowNP) and two replicates, for a total of 2 tissues x 4 treatments x 2 replicates = 16 samples. Libraries were prepared by the Laboratorio de Servicios Genomicos, LANGEBIO, Mexico. www.langebio.cinvestav.mx/labsergen/). Libraries were prepared using the TruSeq RNA Sample Prep Kit v2 (https://support.illumina.com/sequencing/sequencing_kits/truseq_rna_sample_prep_kit_v2.html) and sequenced using the Illumina HiSeq4000 platform at the Vincent J. Coates Genomics Sequencing Laboratory at UC Berkeley, supported by NIH S10 OD018174 Instrumentation Grant, and at Labsergen on the Illumina NextSeq 550 equipment. Transcriptome data are available in the NCBI Sequence Read Archive under study SRP287300 at https://trace.ncbi.nlm.nih.gov/Traces/sra/?study=SRP287300

RNA sequencing reads were aligned against the AGPV3.30 maize gene model set available at Ensembl Plants [111] using kallisto version 0.43.1 [112]).Transcript-level abundance data was pre-processed using R/tximport [113] and summarized at the gene-level before further analysis. Count data were analyzed using a linear model approach in edgeR [114, 115]. We fitted the complete model *counts* ∼ *intercept* + *tissue* * *N-level* * *P-level + error* across the 16 samples. We selected genes-of-interest based on evidence of a non-zero coefficient for at least one model term containing *N-level* or *P-level* (the coef argument to R/edgeR::glmQLFTest included all model coefficients except for the intercept and *tissue* main effect; adjusted FDR < 0.01; absolute log fold change (LFC) > 1; log counts per million (CPM) > 1). An additional subset of 81 NxP interaction genes was selected based on the coefficients *N-level* x *P-level* and *tissue* x *N-level* x *P-level* (adjusted FDR < 0.05; |LFC| >1; logCPM > 1). Genes-of-interest were further categorized based on pairwise LFC for each stress treatment with respect to the full nutrient control for either root or leaves. LFC for each tissue was extracted from the model *counts* ∼ *treatment* + *error*, a threshold of +1 and −1 being used for up- and down-regulation, respectively. Gene functional annotations were assigned as the functional annotation of the blastp reciprocal best hits versus Araport11 [https://doi.org/10.1111/tpj.13415] and uniprot proteins, and the description from the PANNZER2 [https://doi.org/10.1093/nar/gky350] functional annotation webserve. Upset diagrams were generated using R/UpSetR and R/ComplexHeatmap [116, 117]. GO analysis was performed using BiNGO 3.0.3 [118] in the Cytoscape 3.7.2 environment [119] using a hypergeometric test, Benjamini & Hochberg FDR correction and a significance level of 0.05. The Gene ontology file (go.obo) was retrieved from the gene ontology web page (http://geneontology.org/docs/download-ontology/). For each GO category, the mean LFC of the associated genes-of-interest was calculated with respect to each tissue/treatment combination using the pairwise values described above.

### Real-time PCR

For real-time PCR transcript quantification, leaves and roots of five biological replicates per treatment were analyzed. Total RNA was extracted using Trizol and cDNAs were synthesized using SuperScript® II Reverse Transcriptase from Invitrogen (Cat No. 18064071). RT-PCR was performed using 96 well plates in a LightCycler® 480 Instrument by Roche. PCR reactions were performed using KAPA SYBR FAST qPCR Master Mix kit by Kapa Biosystems, with the following cycling conditions: 95 °C for 7 min, followed by 40 cycles of 95 °C for 15 seg; 60 °C for 20 seg; 72°C for 20 seg. The final reaction volume was 10 µl including 1 µl of each 5 µM primer, 1 µl of (40 ng/µl) template cDNA, 5 µl of SYBR Master Mix and 2 µl of distilled water. The relative quantification of the gene expression was determined as 2^ΔCt^, where ΔCt = 2^(Average Ct of reference genes - Ct of gene of interest) [120]. Values reported are the mean of five biological replicas ± SE of one representative experiment. Previously described reference genes [121] were used as controls: Cyclin-Dependent Kinase (*Cdk*; GRMZM2G149286) and a gene encoding an uncharacterized protein (*Unknown*; GRMZM2G047204). PCR primers were designed using Primer3Plus software [122] and are listed in MZ95_RT_Primers in Supplemental File 3.

### Phylogenetic analysis of the SPX-domain protein family

Maize putative SPX-domain protein encoding genes were identified using a methodology previously described for the maize *Pap* gene family [58]. Briefly, *Arabidopsis* and rice proteins [64] were retrieved and aligned using MUSCLE v3.8 [123]. The alignment was then converted to Stockholm format. B73 maize primary transcript predicted protein sequences v3.31 [124] obtained from Ensembl Plants [108] were searched using HMMER suite version 3.1b2 [125]. After manually checking and filtering for proteins lacking the canonical SPX domain [126], 15 putative SPX-protein sequences were identified. Where noted, gene models annotated in the v4 genome assembly were preferred. For phylogenetic analysis, *Arabidopsis*, rice and maize SPX proteins were aligned using MUSCLE [123] and passed to MEGA version X [127, 128]. We manually selected SPX sub-domains defined by [64] and corrected some mismatches in the alignment (Fig. S3). A 1000 bootstrap phylogenetic tree was constructed with Maximum Likelihood method and Le_Gascuel_2008 model [129].

## ABBREVIATIONS

DAE: Days After Emergence
FDR: False Discovery Rate
GO: Gene Ontology
ICP-MS: Inductively Coupled Plasma Mass Spectrometry
LFC: Log_2_ Fold Change
NSR: Nitrogen Starvation Response
PCR: Polymerase Chain Reaction
PSR: Phosphate Starvation Response

## DECLARATIONS

### Ethics approval and consent to participate

Not applicable

### Consent for publication

Not applicable

### Availability of data and materials

Transcriptome data are available in the NCBI Sequence Read Archive under study SRP287300 at https://trace.ncbi.nlm.nih.gov/Traces/sra/?study=SRP287300

### Author Contributions

JVT-R and MNS-V conceived and designed the experiments, performed the experiments, analyzed the data, prepared figures and tables and authored drafts of the paper.

RACM analyzed RNA-seq data and authored drafts of the paper.

JAM-S performed experiments, analyzed the data, and authored drafts of the paper.

CSG conceived and designed the experiments, contributed reagents/materials/analysis tools and authored or reviewed drafts of the paper.

RJHS conceived and designed the experiments, analyzed the data, contributed reagents/materials/analysis tools, prepared figures and/or tables, authored or reviewed drafts of the paper.

All authors approved the final draft.

## Acknowledgements

We thank Benjamin Barrales-Gamez, Ana Laura Alonso-Nieves and Jessica Carcaño-Macias for assistance in plant growth and harvest.

## ADDITIONAL FILES

Figure S1. Growth traits for plants grown under Full, LowN, LowP and LowNP. Plots show estimated coefficient and associated standard error. The significance of the treatment effect is shown as *** p <0.001, ** p <0.01, * p <0.05, p<0.1 (Kruskal-Wallis test; p-value adjusted for multiple tests). Lowercase letters indicate significant (p<0.05) pairwise differences (Dunn test). Figure accompanies MZ66_Growth_Analysis in Supplemental File 1.

Figure S2. Endpoint traits for plants grown under Full, LowN, LowP and LowNP. Plots show estimated coefficient and associated standard error. The significance of the treatment effect is shown as *** p <0.001, ** p <0.01, * p <0.05, p<0.1 (Kruskal-Wallis test; p-value adjusted for multiple tests). Lowercase letters indicate significant (p<0.05) pairwise differences (Dunn test). Figure accompanies MZ66_Endpoint_Analysis in Supplemental File 1.

Figure S3. GiaRoot root features for plants grown under Full, LowN, LowP and LowNP. Plots show estimated coefficient and associated standard error. The significance of the treatment effect is shown as *** p <0.001, ** p <0.01, * p <0.05, p<0.1 (Kruskal-Wallis test; p-value adjusted for multiple tests). Lowercase letters indicate significant (p<0.05) pairwise differences (Dunn test). Figure accompanies MZ66_Giaroots_Analysis in Supplemental File 1.

Figure S4. Ions concentrations for plants grown under Full, LowN, LowP and LowNP. Plots show estimated coefficient and associated standard error. The significance of the treatment effect is shown as *** p <0.001, ** p <0.01, * p <0.05, p<0.1 (ANOVA; p-value adjusted for multiple tests). Lowercase letters indicate significant (p<0.05) pairwise differences (Tukey). Figure accompanies MZ66_Ionomics_Analysis in Supplemental File 1.

Figure S5. Transcription of PSR is reduced under LowN availability. Scatter plot showing the distribution of transcript accumulation (log_2_ fold change, LFC) of 1, 555 genes in A) leaves and B) roots in LowP and lowN. Dotted lines represent LFC of −1 and 1. Dots filled using heat-colors showing LowNP transcript accumulation. C) Differential transcript accumulation (z, row standardized LFC) with respect to Full of the top 30 (ranked by FDR) classic genes. Figure accompanies MZ67_Selected_Classics in Supplemental File 2.

Figure S6. SPX-domain family members respond to reduced N and P availability. A) Phylogenetic tree of SPX-domain family proteins in maize. Likelihood tree built with Arabidopsis, rice and maize SPX-domain proteins. Numbers at the nodes indicate bootstrap (1000) support as percentage. B) Heat map of maize SPX-domain gene family expression under LowN, LowP and combined LowNP with respect to Full (z, row standardized log_2_ fold change). Asterisks indicate genes identified as regulated in the transcriptome analysis.

Supplemental file 1. Plant growth to 40 days after emergence (experiment MZ66). Workbook contains raw and processed phenotypic data from maize plants under lowN, lowP and lowNP until 40 days after emergence.

Supplemental file 2. Transcriptome analysis of from maize plants under lowN, lowP and lowNP until 25 days after emergence (experiment MZ67). Workbook contains count data, analysis, candidate gene lists and annotation, and enriched GO terms.

Supplemental file 3. MZ95 experiment. File contains phenotypic and gene expression data from qRT-PCR of maize leaves and roots under LowN, LowP and lowNP at 25 days after emergence. Workbook contains expression data and analysis.

