## Supplementary figures and images for "Low nitrogen availability inhibits the phosphorus starvation response in maize (*Zea mays* ssp. *mays* L.)"

### Figure S1

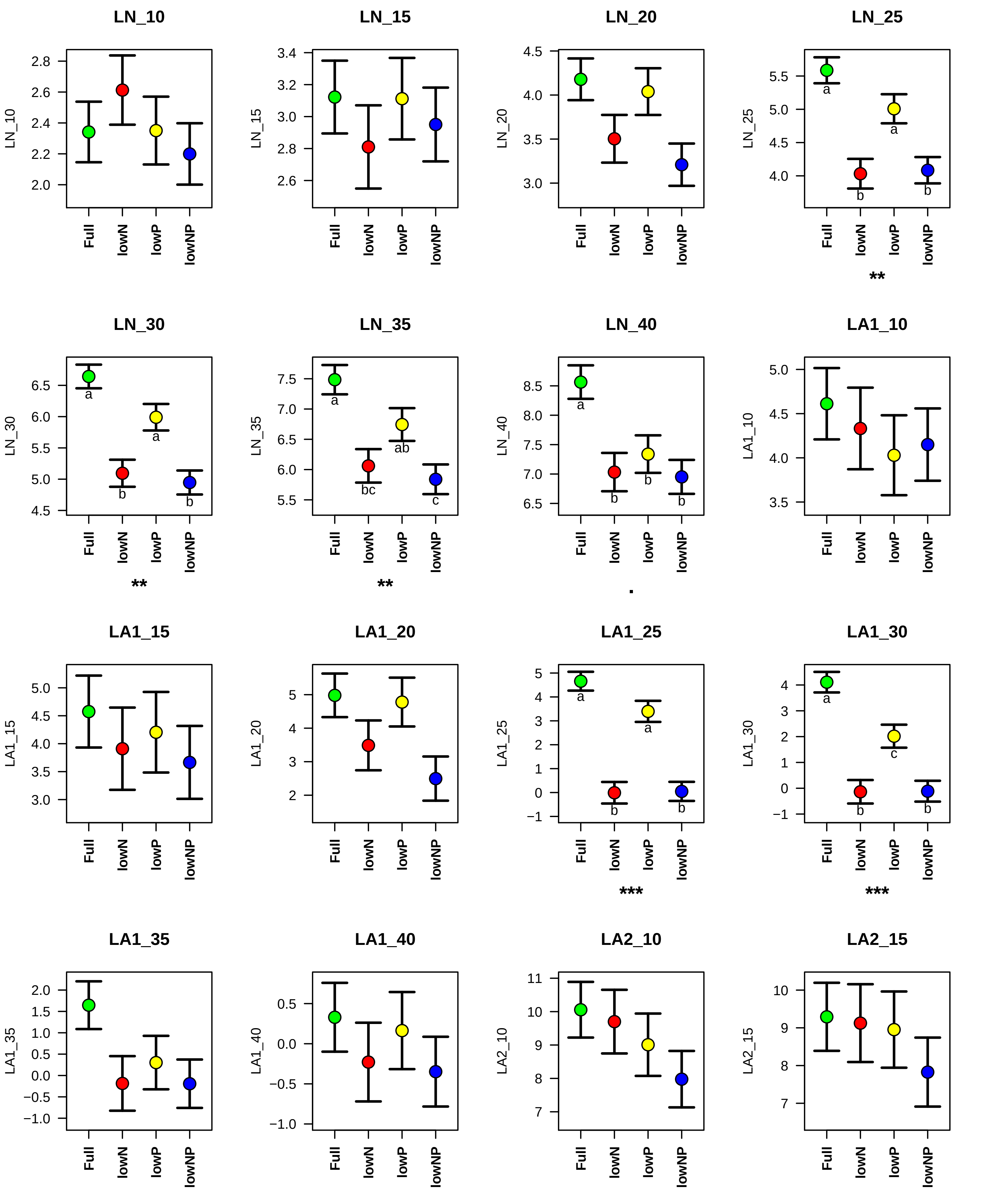

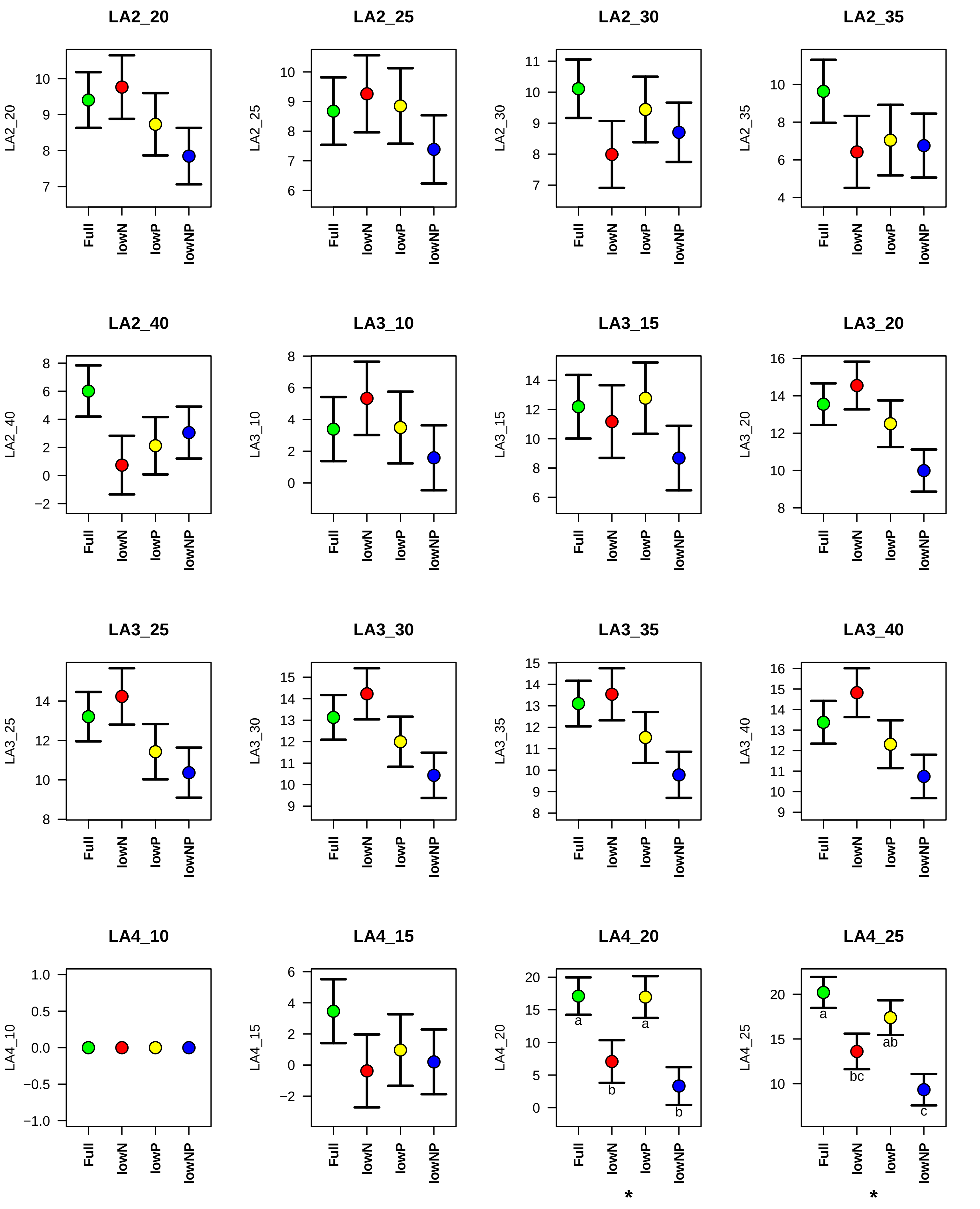

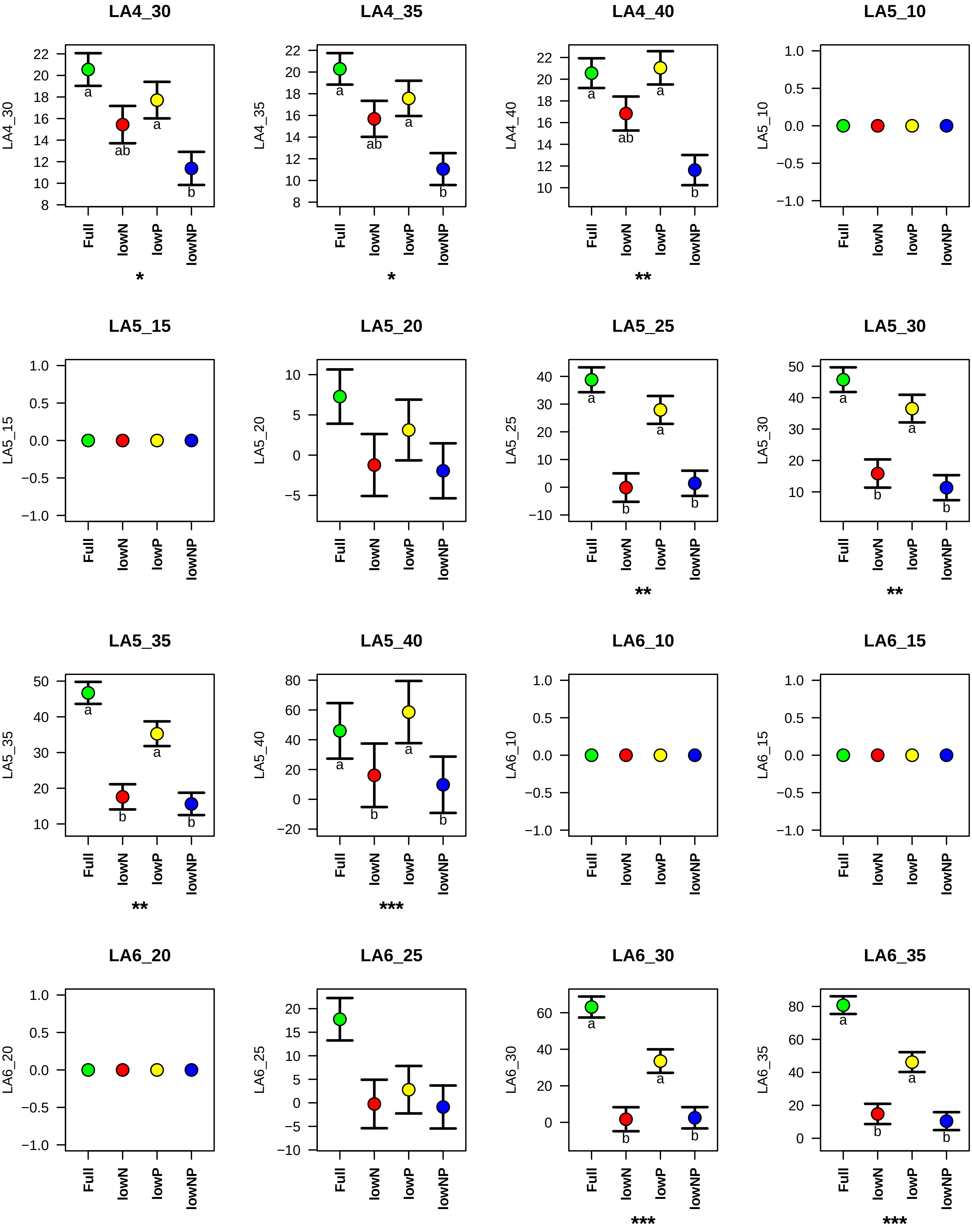

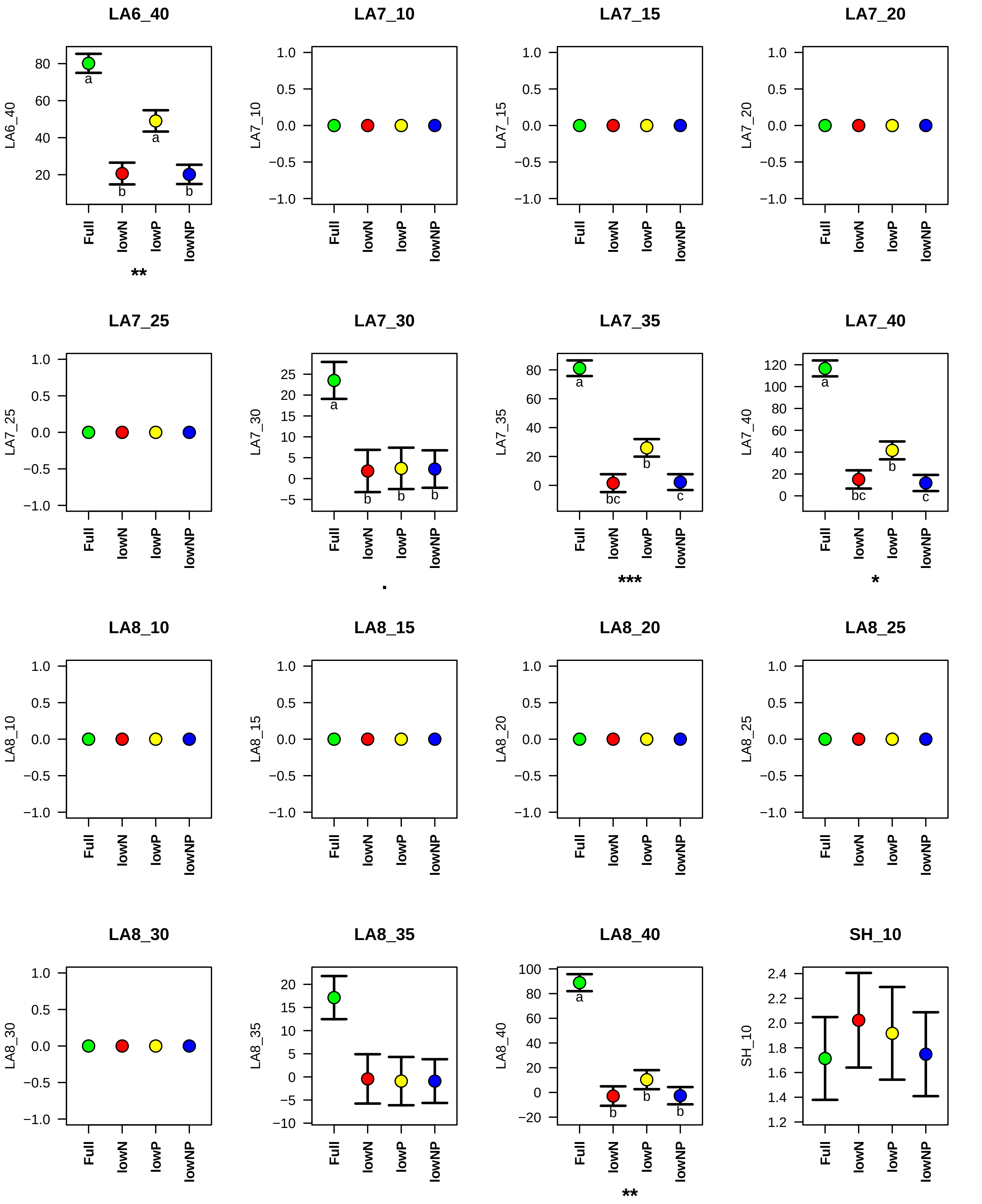

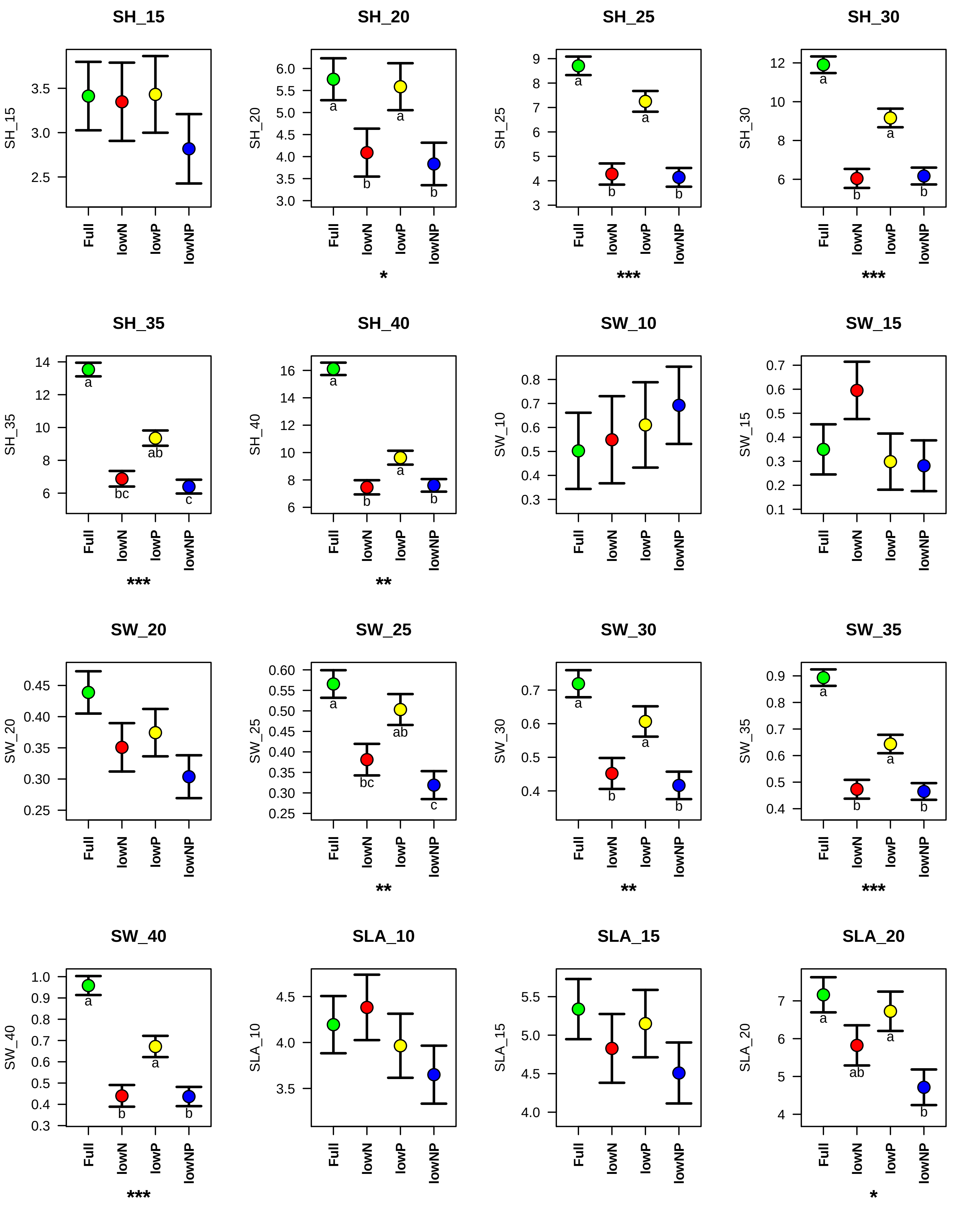

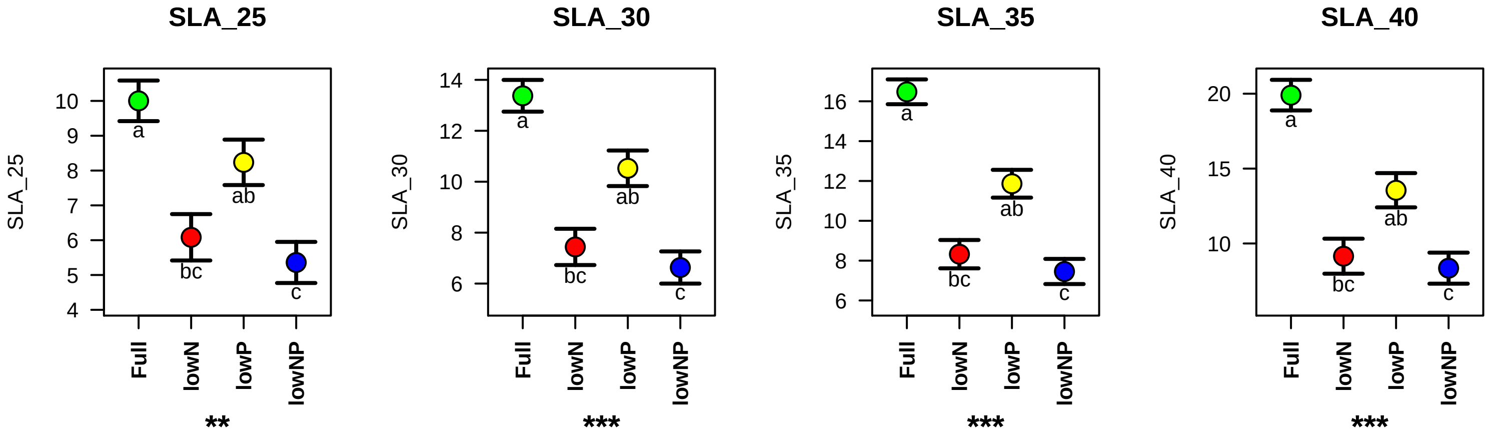

### Figure S2

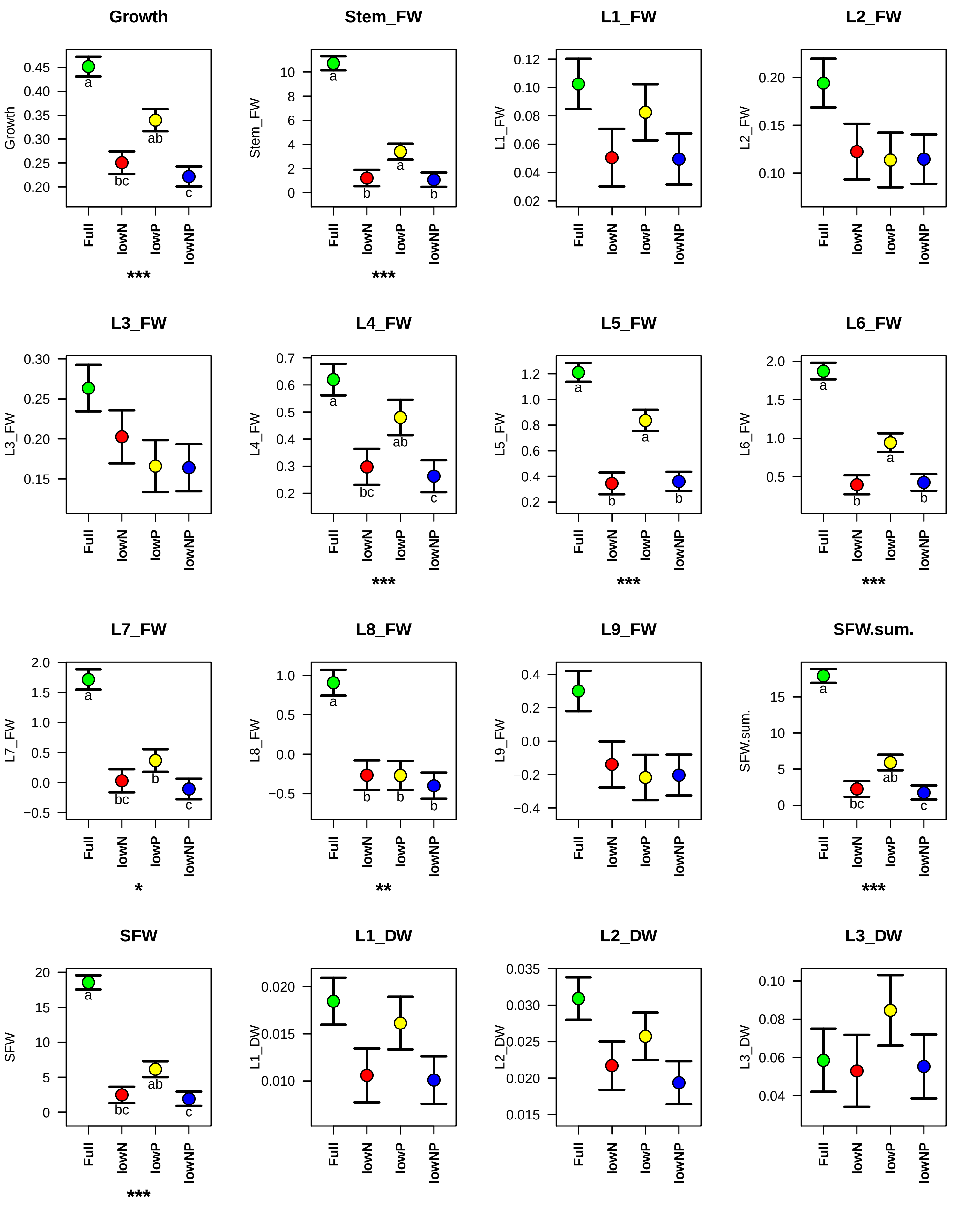

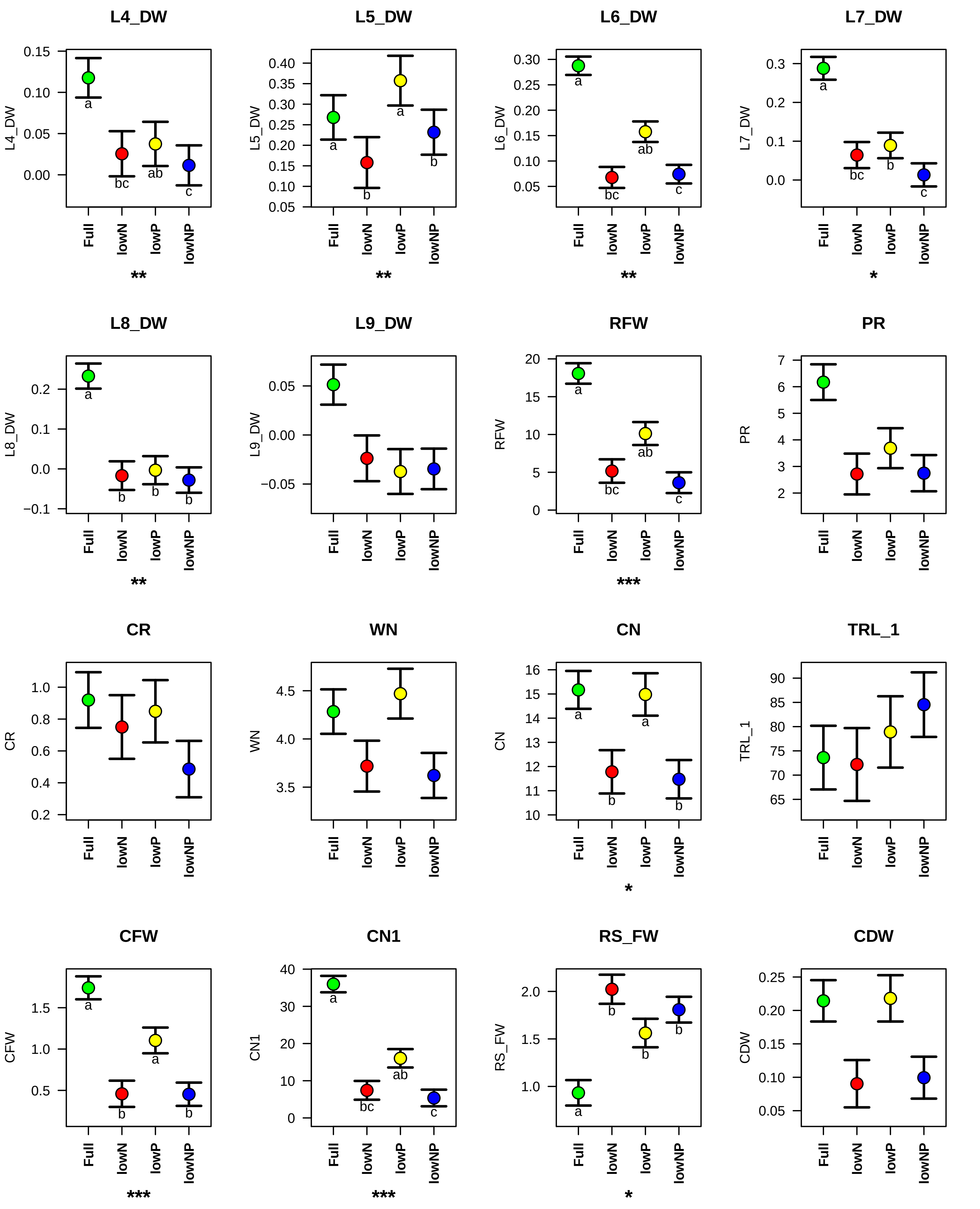

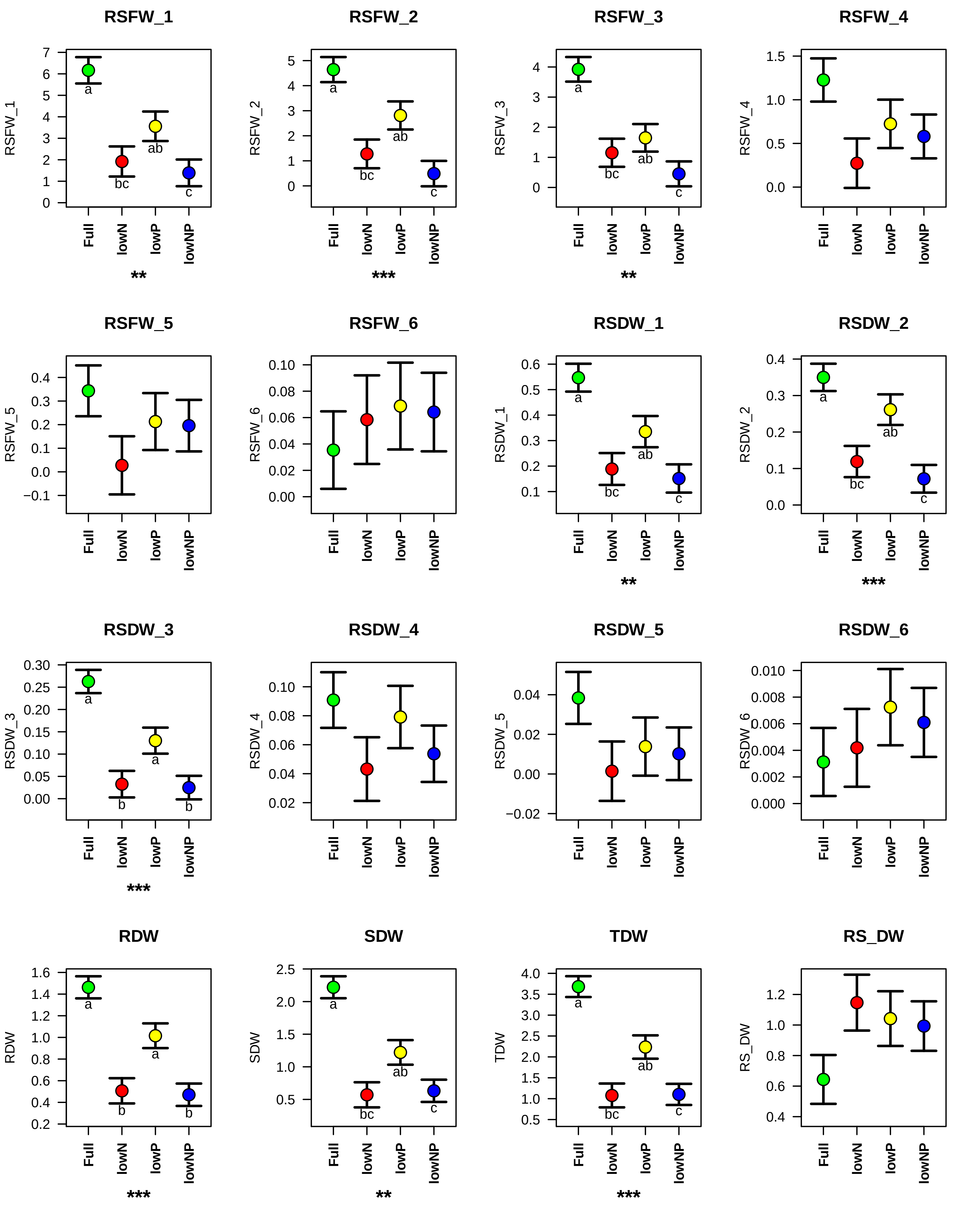

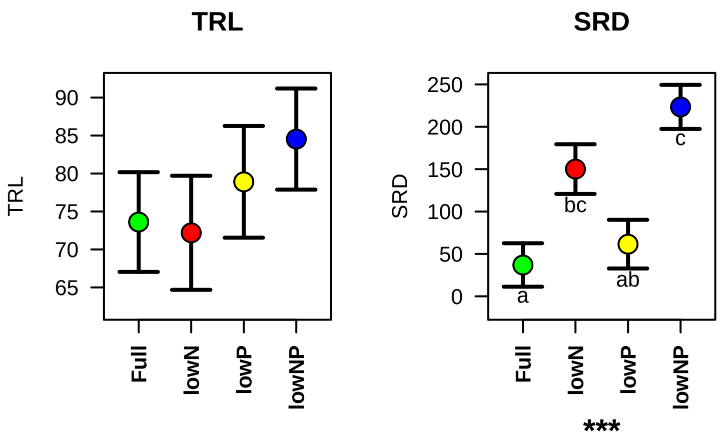

### Figure S3

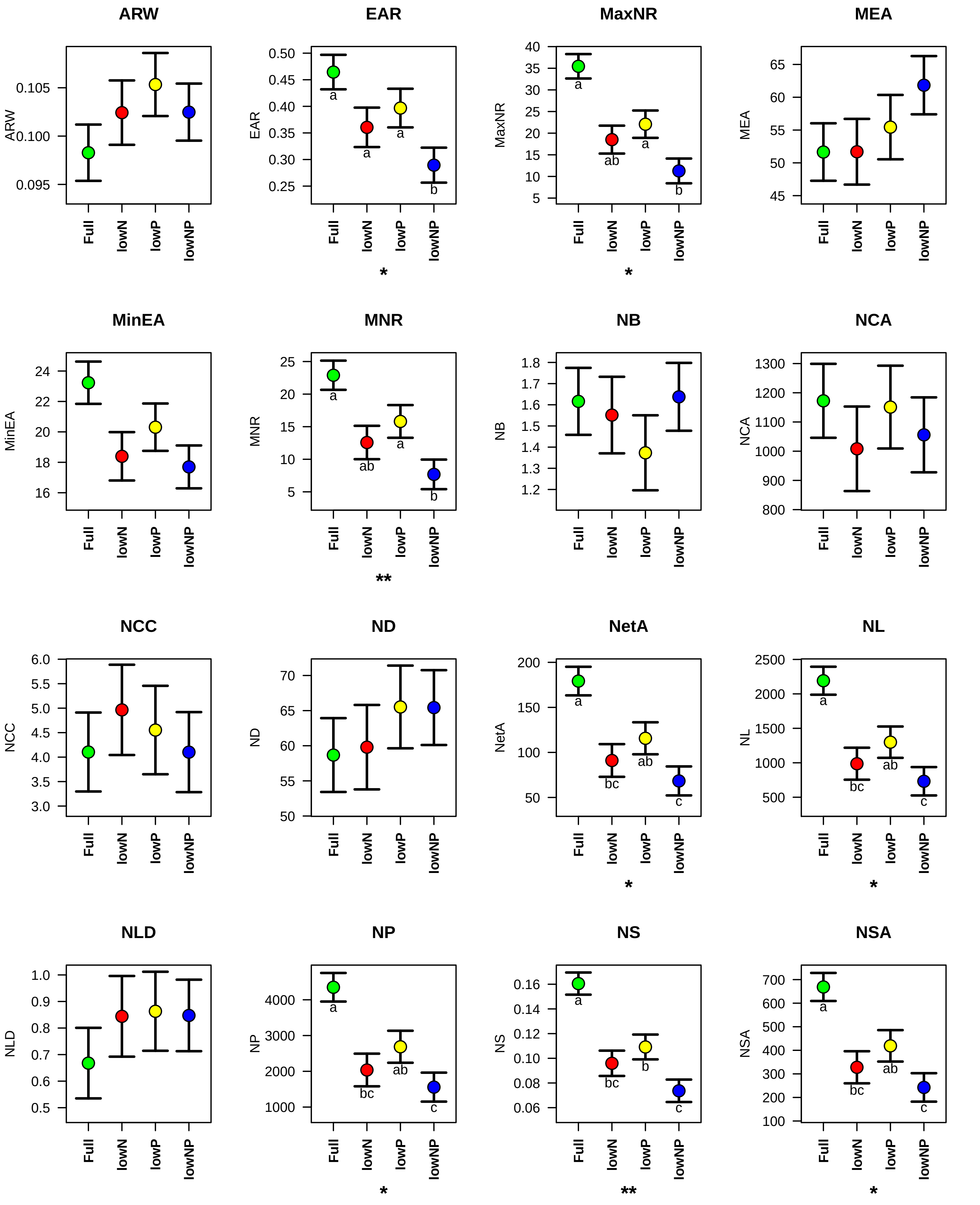

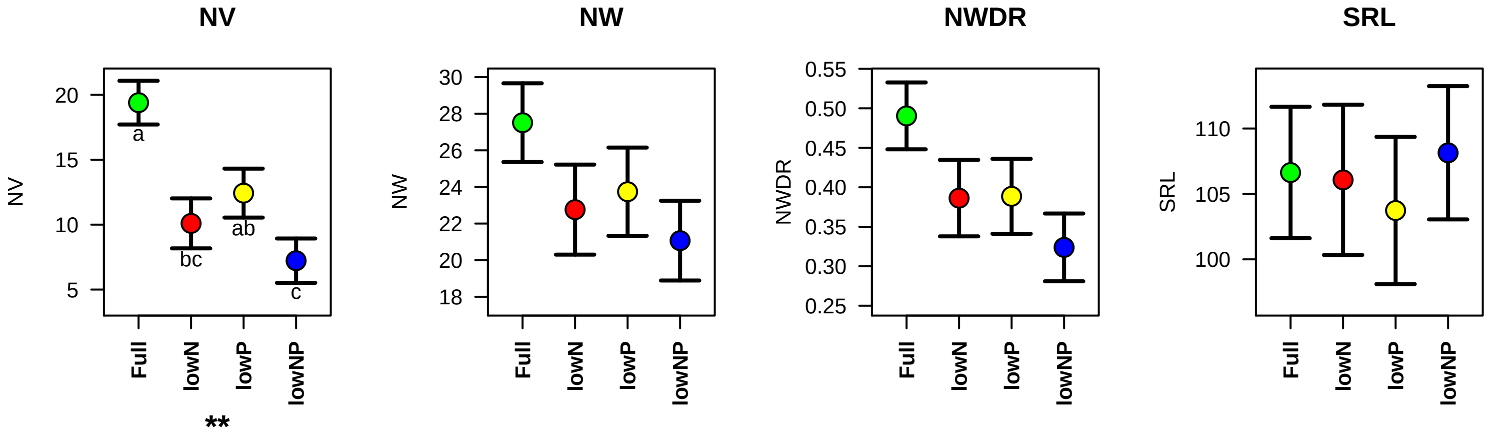

### Figure S4

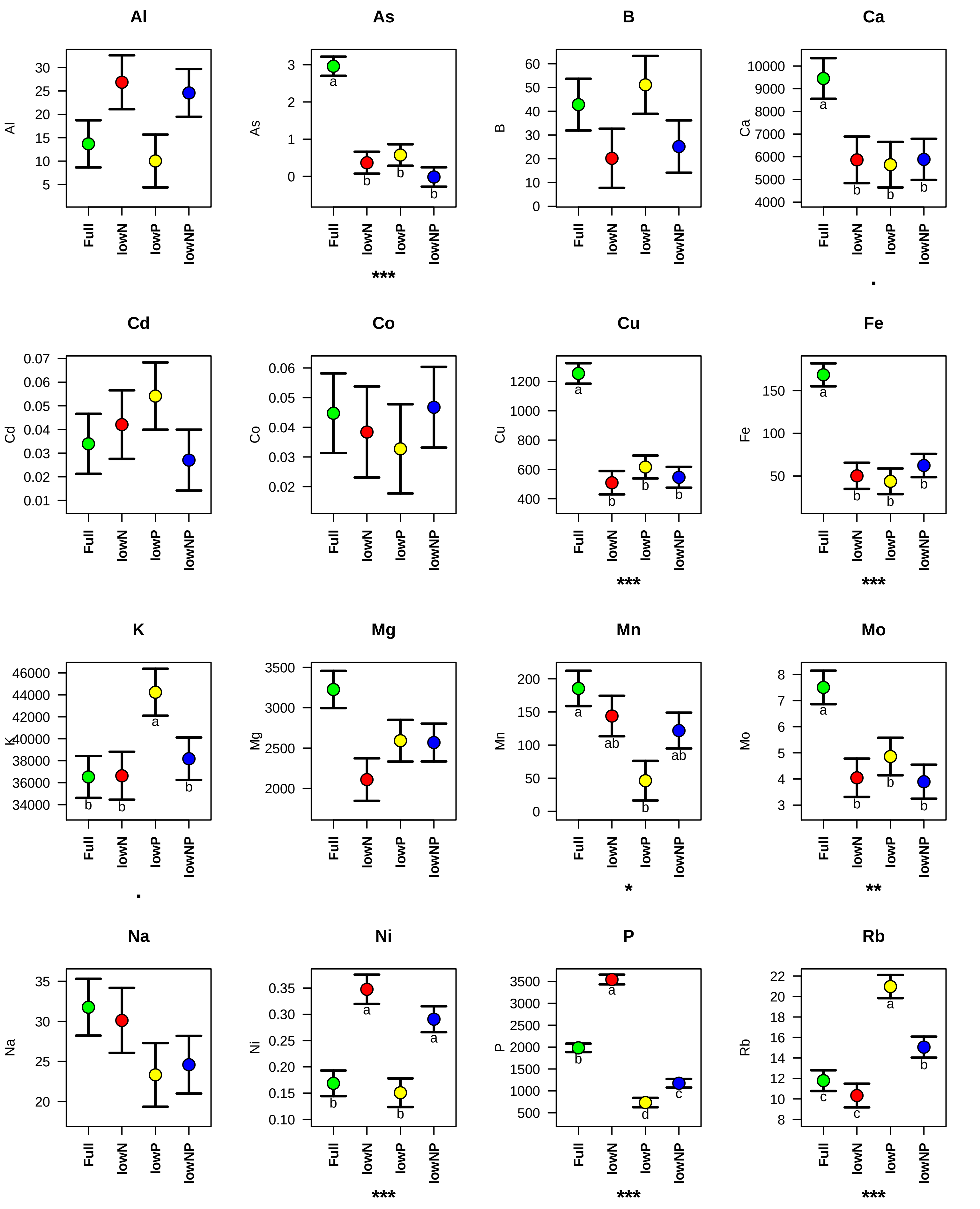

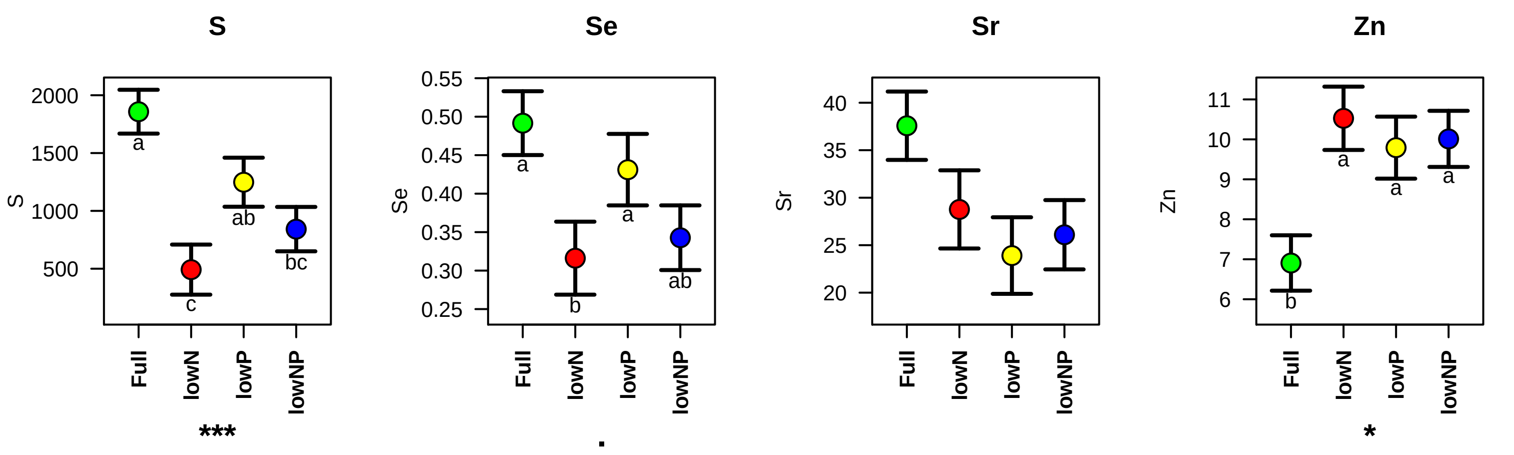

### Figure S5

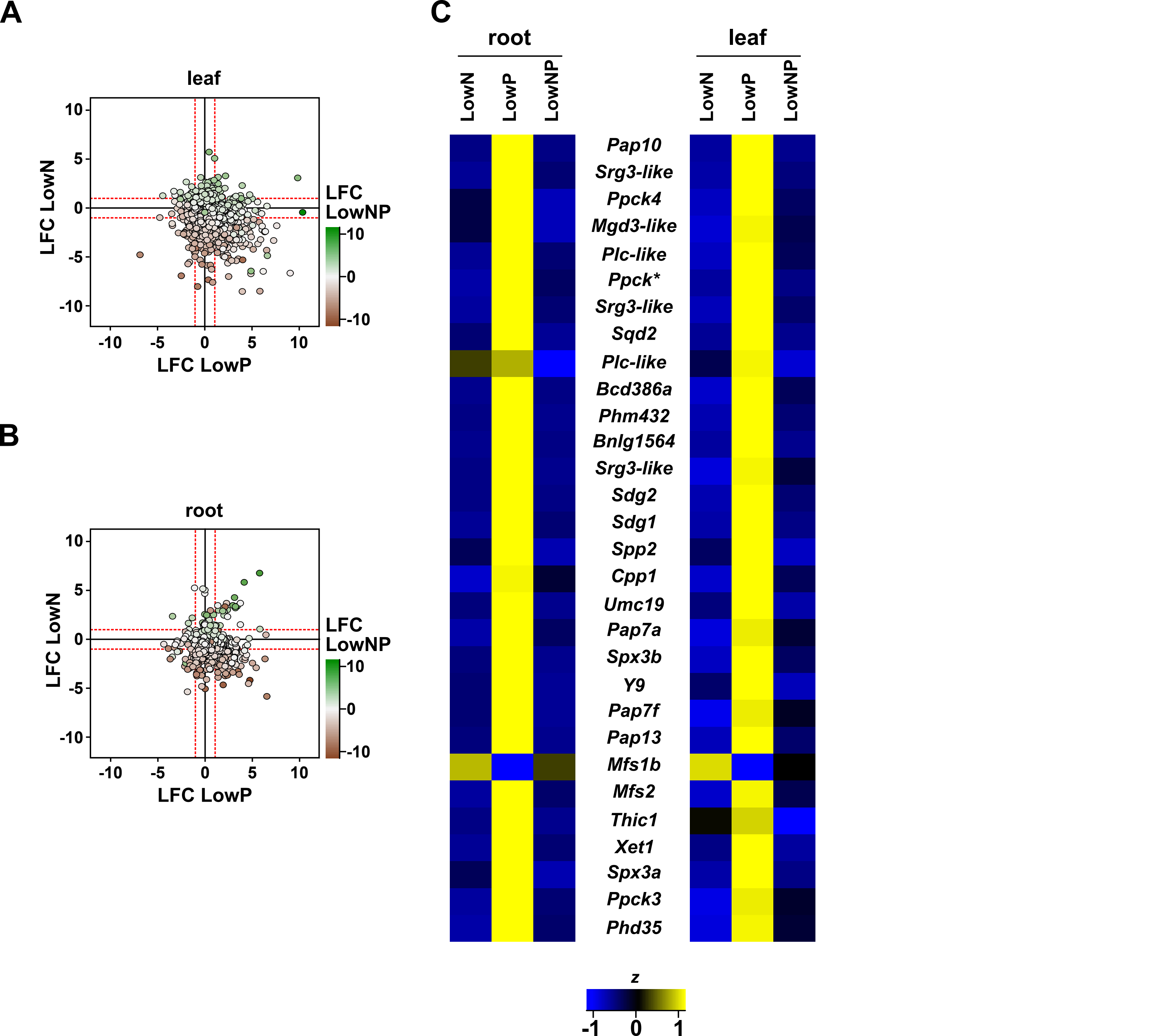

### Figure S6

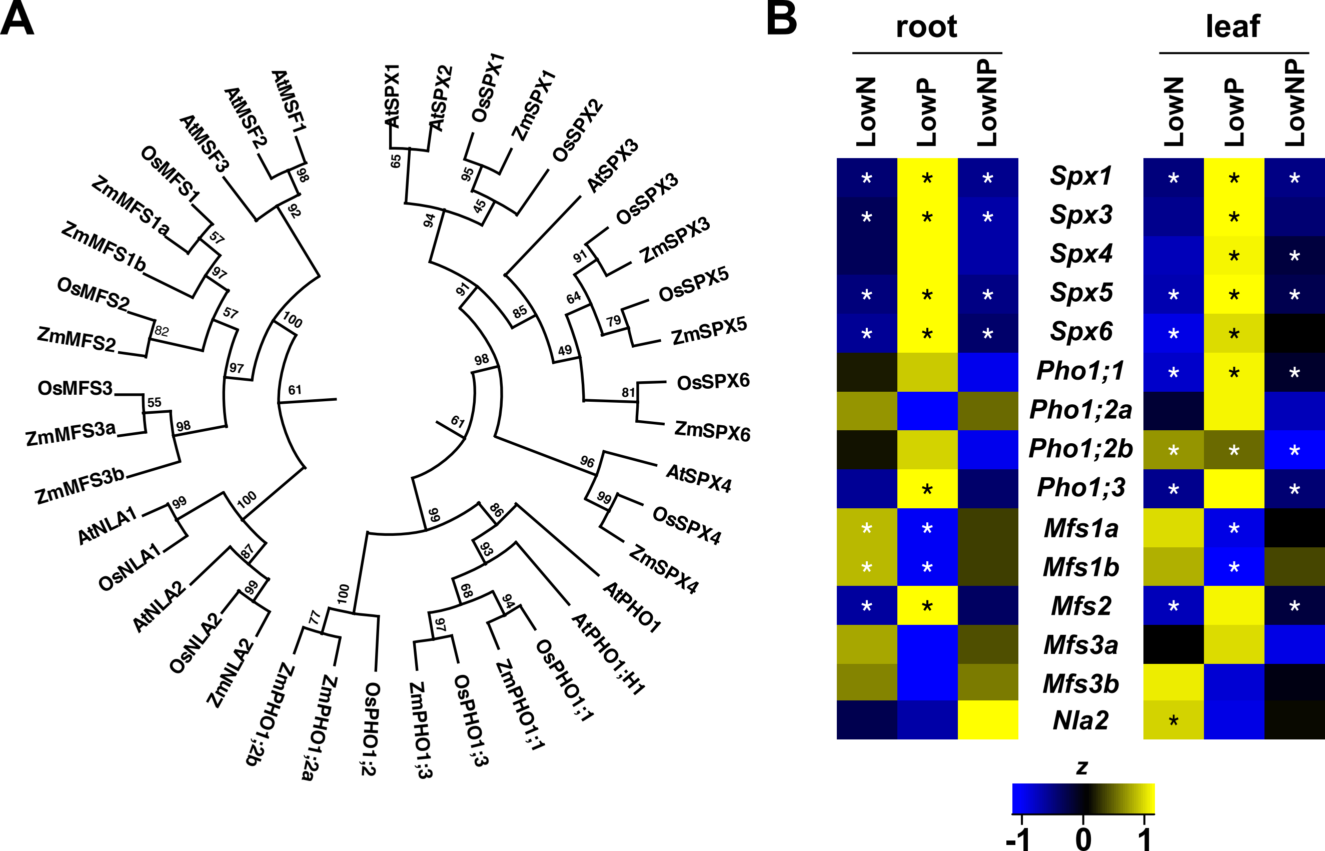
